# PepGlider: Property Regularized VAE for interpretable and controllable peptide design

**DOI:** 10.1101/2025.10.13.682099

**Authors:** Aleksandra Gwiazda, Paulina Szymczak, Ewa Szczurek

## Abstract

Computational peptide design requires precise control over peptide properties that often exhibit complex correlations. Existing generative models for peptide design rely on simplistic discrete conditioning mechanisms rather than precise targeting of specific property values. We present PepGlider, a continuous property regularization framework that enables direct control over their specific values. The method achieves structured latent space with superior disentanglement quality and displays smooth property gradients along regularized dimension. *In silico* experimental results demonstrate that PepGlider enables independent control of naturally correlated properties, and supports both *de novo* generation and targeted optimization of existing peptides. PepGlider applied to antimicrobial peptide design allows generation of candidates with desired antibacterial activity profile and maintained low toxicity profile. Unlike existing approaches, PepGlider provides precise control over continuous property distributions while maintaining generation quality, thus offering a generalizable solution for therapeutic and materials applications requiring exact property specifications.

## 1 Introduction

Peptide design across diverse biomedical applications confronts a fundamental optimization challenge: achieving precise control over continuous peptide properties that often exhibit complex correlations or direct conflicts. Antimicrobial peptide (AMP) design has emerged as particularly urgent due to the escalating antimicrobial resistance crisis. Multidrug-resistant pathogens cause over 700,000 deaths annually, with projections reaching 10 million by 2050 without intervention (O’Neill, 2016). AMPs are a promising alternative with broad-spectrum activity, rapid bacterial killing kinetics, and reduced resistance development compared to conventional antibiotics (Hancock and Sahl, 2006). The need for controllable peptide design is particularly evident in AMPs, where antimicrobial efficacy depends on complex interplay between physicochemical properties such as charge or hydrophobicity (Hancock and Sahl, 2006). Introducing positively charged amino acids to increase net charge - crucial for membrane interaction - often disrupts the distribution of hydrophobic residues essential for bacterial killing. Such natural properties correlation complicate the optimization of antimicrobial activity, exemplifying the broader challenge of achieving precise control of them.

Deep generative models have emerged as powerful tools for peptide sequence design, but current approaches exhibit significant limitations in controllability and precision (Szymczak and Szczurek, 2023). Existing conditional generation frameworks rely predominantly on discrete conditioning mechanisms that fail to enable precise targeting of specific continuous property values. This limits optimization to coarse-grained categories rather than exact property ranges required for functional applications. Recent advances in attribute-controllable generation, particularly AR-VAE (Pati and Lerch, 2021), structure latent spaces such that specific dimensions encode target properties through monotonic relationships. However, AR-VAE’s discrete signum-based regularization of the loss function only enable relative ordering between samples, not precise targeting of specific property values. This limitation renders AR-VAE unsuitable for peptide design applications.

To address these challenges in controllable peptide design, we present PepGlider, a continuous property regularization framework that achieves precise, independent control over correlated peptide properties. Figure 1a illustrates our framework, which supports two generation modes: unconstrained generation for *de novo* peptide discovery through targeted sampling from the prior distribution, and analog generation for optimizing existing peptides via targeted latent manipulations. Our approach makes three main technical contributions:

**Figure 1.**
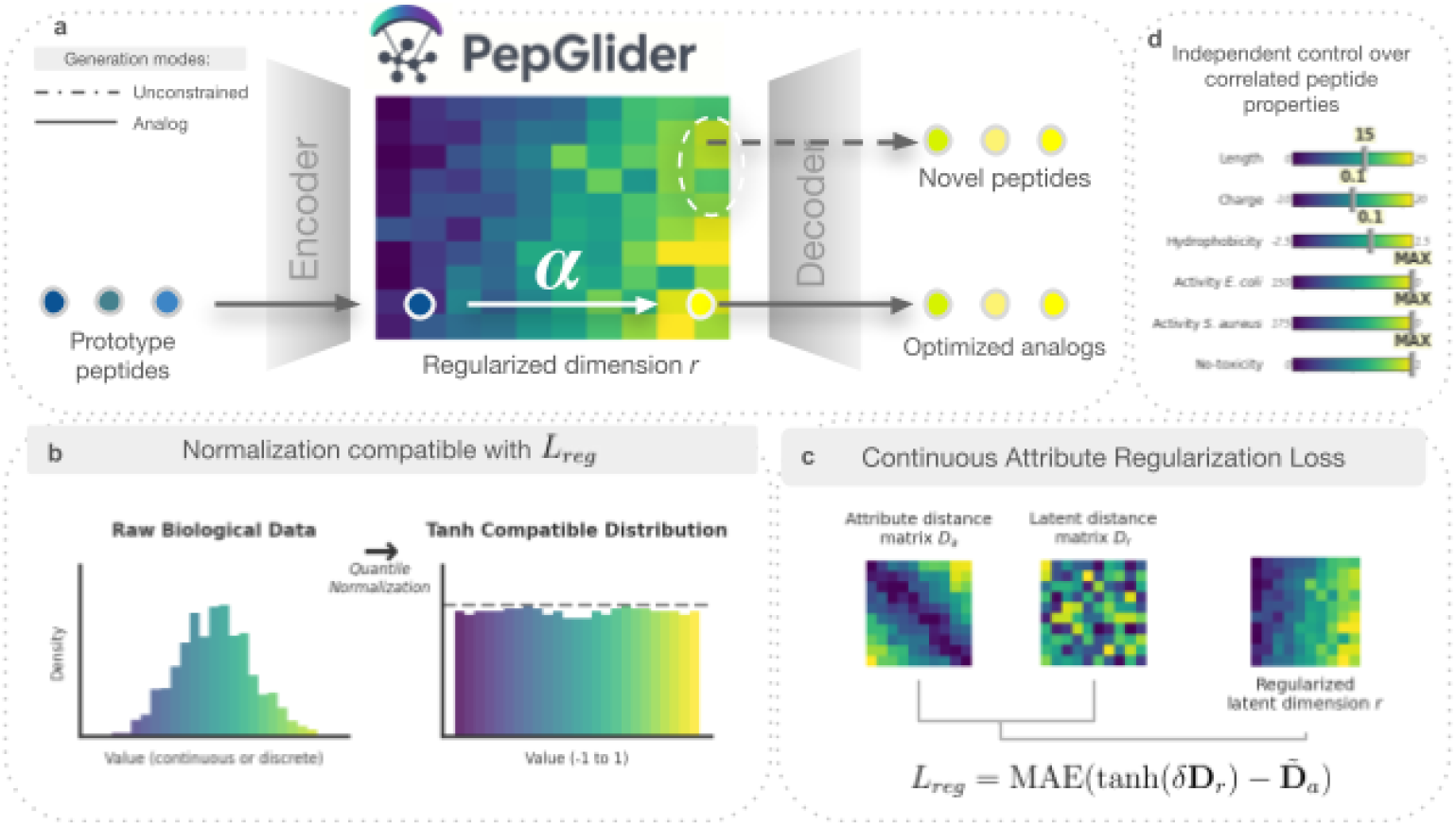
PepGlider: Continuous property regularization for controlled peptide design. **(a)** PepGlider overview showing unconstrained generation through targeted sampling from the prior distribution and analog generation modes, where prototype peptides are encoded, modified via displacement values *α* in regularized latent dimensions, and decoded to generate optimized peptides. **(b)** Normalization procedure scaling raw property values to [−1, 1] range to match *tanh* activation for continuous loss optimization. **(c)** Continuous property regularization loss aligning attribute distance matrix *D*_*a*_ with latent distance matrix *D*_*r*_ through MAE loss. **(d)** Independent control over correlated peptide properties (length, charge, hydrophobicity, activity, non-toxicity) enabled by structured latent space.

- (i) we introduce property-specific normalization strategies (Figure 1b) that scale diverse peptide properties to a unified range while preserving biological relevance, particularly adaptive range normalization for clinically relevant antimicrobial potency ranges
- (ii) we extend AR-VAE with continuous loss formulations (Figure 1c) that enable precise targeting of specific peptide property values rather than relative orderings, and
- (iii) we demonstrate independent manipulation of naturally correlated peptide properties through structured latent space design (Figure 1d), enabling multi-objective optimization across correlated properties.

The resulting framework offers a generalizable methodology for precise property control across diverse peptide design applications.

## 2 Related work

Controllable peptide design intersects multiple research areas, including conditional generation and latent space regularization, each addressing different aspects of the challenge of navigating correlated peptide properties.

### Controllable Peptide Design

Current approaches to controllable peptide generation, particularly for AMPs, employ three main strategies: conditional generation, post-hoc filtering, and guidance during sampling. While conditional methods like HydrAMP (Szymczak et al., 2023) directly incorporate constraints, they are limited to binary classification. Post-hoc approaches (Das et al., 2021; Pandi et al., 2023; Torres et al., 2025) suffer from severe efficiency limitations when targeting rare property combinations. The exponential search space of peptide sequences makes exhaustive sampling impractical, particularly when multiple properties must be optimized simultaneously. Guidance-based methods attempt to steer the generation process toward desired properties during sampling, including approaches that use Monte Carlo Tree Guidance (Tang et al., 2025) and reinforcement learning with property-based rewards (Wang et al., 2024). However, guidance approaches operate at the sampling level rather than embedding controllability into the learned representation, making them computationally expensive during generation and unable to leverage the structured relationships between properties for more efficient optimization.

### Latent Space Regularization

Latent space regularization methods from other domains learn representations where properties naturally align with latent structure. VAE-based approaches have pioneered this direction through various regularization strategies. CorrVAE (Wang et al., 2022) addresses property correlations through specialized loss functions designed to handle interdependent data characteristics. Property-controllable VAE (Guo et al., 2020) incorporates property prediction losses directly into the variational objective, creating latent representations that encode desired features. Conditional Subspace VAE (Klys et al., 2018) partitions the latent space according to property-specific regions, enabling targeted sampling from relevant subspaces. AR-VAE (Pati and Lerch, 2021) aligns latent and property spaces through distance matrix matching to create monotonic relationships between latent dimensions and target properties. While these latent space methods offer promising frameworks for controllable generation, they have not been adapted to address the specific challenges of peptide design, particularly the need for precise property targeting across correlated physicochemical characteristics and efficient access to rare, but functional attribute combinations essential for therapeutic applications.

## 3 Methods

### 3.1 Background

We build PepGlider upon the Attribute-Regularized VAE framework (Pati and Lerch, 2021), which structures classic VAE latent representations such that specific dimensions encode target attributes in a monotonic fashion. Let **x** denote a data sample (here, a peptide sequence) and **z** represent the corresponding latent representation obtained through the VAE encoder. For each mini-batch of size *m*, the method constructs property and latent distance matrices for all pairs of samples *i, j* ∈ {1, …, *m*}:

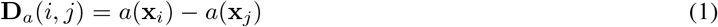

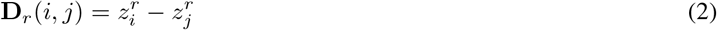

where *a*(*·*) represents the attribute function, *r* denotes the regularized latent dimension, **D**_*a*_ is the attribute distance matrix, and **D**_*r*_ is the latent distance matrix. The regularization loss enforces alignment between these distance matrices:

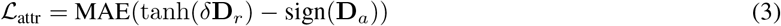

The complete AR-VAE objective combines this with standard VAE components (Kingma and Welling, 2013):

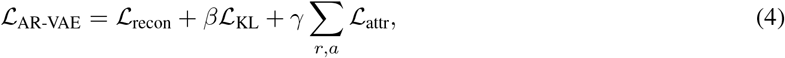

where *β* controls the weight of the Kullback-Leibler regularization term, the parameter *δ* controls the spread of latent vectors, and *γ* weights the overall attribute-based regularization strength.

### 3.2 PepGlider Framework

PepGlider is designed for general peptide design applications requiring precise property control across diverse peptide optimization objectives. The signum function in AR-VAE creates discrete comparisons that limit controllability to relative ordering rather than absolute values. We address this limitation through two key innovations:

- **Continuous Property-Based Regularization** To overcome inherent AR-VAE limitation, we introduce continuous property-based regularization. In the continuous formulation of the loss function we discard the signum-based comparison, and introduce the property regularization term:

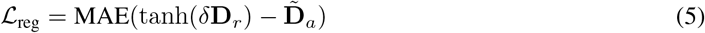

The modified PepGlider loss becomes:

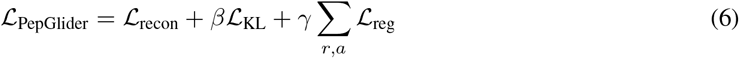
- **Property Normalization** To enable compatibility with our continuous loss formulation, we introduce property-specific normalization strategies that scale 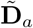 to [−1, 1] while preserving biological meaning.

For properties with symmetric importance across their range, we apply **quantile transformation** (QT). Raw values are transformed via quantile transformation *Q*(*·*) to *U* (0, 1), then linearly scaled to [−1, 1]:

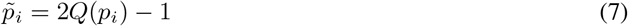

This ensures uniform property space coverage and eliminates scale bias.

For properties where specific value ranges hold disproportionate biological importance, we employ **adaptive range normalization** that allocates normalized space non-uniformly. This approach dedicates greater resolution to critical regions while compressing less important ranges. Values are mapped through region-specific empirical CDFs. Detailed equations and specific applications are provided in Appendix A.1.4.

This normalization framework, combined with continuous property-based regularization, ensures all properties and latent regularization terms operate within the same bounded range, enabling PepGlider to achieve precise, quantitative control over multiple peptide properties simultaneously.

#### 3.2.1 Controlled Peptide Generation Modes

Post-training, PepGlider enables two generation modes that leverage the organized latent space for different peptide design objectives:

- **Analog generation** modifies existing peptides through manipulation of regularized dimensions. Given input sequence **x** with latent code **z** = Enc(**x**), property *a* associated with regularized dimension *r* is controlled by displacement: 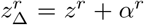, with 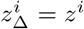 for all *i* ≠ *r*. The displacement *α*^*r*^ is determined by the target property change, exploiting the monotonic relationship between *z*^*r*^ and *a*(**x**) established during training. The modified sequence is reconstructed as 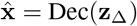. For multi-property control, displacements are applied simultaneously across regularized dimensions *{r*_1_, *r*_2_, …, *r*_*k*_*}*.
- **Unconstrained generation** samples latent codes **z** ∼ *N* (**0, I**) from the prior distribution and decodes them to generate peptides. To generate peptides with target property values, the regularized dimension *r* is set using displacement to a specific value corresponding to the desired property level, while the remaining dimensions are sampled from *N* (0, 1). The generated sequence is obtained as 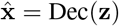.

## 4 Experimental setup

### 4.1 Regularized properties

#### Physicochemical features

PepGlider targets three fundamental physicochemical properties that serve as key determinants of antimicrobial activity: **net charge** (C, calculated at physiological pH), **hydrophobicity** (H, computed as the average hydrophobicity value across all residues in the sequence using established amino acid hydrophobicity scales), and **sequence length** (L). The selection and computational implementation of these properties is described in Appendix A.1.1.

#### Antimicrobial activity

We incorporate Minimum Inhibitory Concentration (MIC) values, the established quantitative measure of antimicrobial potency, as continuous properties predicted by APEX (Wan et al., 2024), a deep learning model trained on experimentally validated antimicrobial activity data. We target two clinically relevant bacterial pathogens: ***Escherichia coli*** (*E. coli*), a Gram-negative bacterium, and ***Staphylococcus aureus*** (*S. aureus*), a Grampositive pathogen, both representing major causes of antibiotic-resistant infections. For each pathogen, MIC predictions are averaged across multiple strains including drug-resistant variants to ensure robust activity estimates. Detailed information on MIC definitions, APEX architecture, and target strains is provided in Appendix A.1.2.

#### Non-toxicity

To enable non-toxicity assessment of generated peptides, we trained a binary classifier to predict **hemotoxicity**. While existing toxicity predictors are available (Rathore et al., 2025; Salem et al., 2022), we developed a custom classifier to maximize utilization of experimental hemolytic activity data from the DBAASP database (Pirtskhalava et al., 2021). Specifically, we extract HC50 measurements (peptide concentration causing 50% hemolysis) and apply rule-based binarization to convert continuous HC50 values into binary toxicity labels. To expand the non-toxic class and improve classifier robustness, we supplemented DBAASP data with metabolic, signal, and hormone peptides from Peptipedia (Cabas-Mora et al., 2024), which are presumed non-hemolytic due to their physiological roles. This combined approach enables incorporation of a broader range of experimental and functional data (detailed procedure in Appendix A.1.3). Peptide sequences were featurized using a comprehensive set of physicochemical property values, comprising over 100 molecular descriptors including basic properties (length, charge, hydrophobicity), structural descriptors (secondary structure fractions, topological features), and specialized amino acid scales. An XGBoost classifier was trained on these features to predict binary non-toxicity (1 = non-toxic, 0 = toxic).

We employ quantile normalization for physicochemical properties (charge, length, hydrophobicity) and non-toxicity predictions, and adaptive range normalization for MIC values, providing granular representation of clinically relevant ranges (0-32 *µ*g/ml) while maintaining the [− 1, 1] scaling required for direct alignment with tanh(*d***D**_*r*_) outputs. Technical implementation details are in Appendix A.1.4, with data distribution before and after normalization shown in Figure S2.

### 4.2 Models

PepGlider implementation details and training procedures are provided in Appendix A.3. We evaluate PepGlider against six baseline approaches that represent different paradigms for controlled peptide generation and enable systematic assessment of our methodological contributions. These include ablation variants to isolate the impact of our key innovations (VAE, AR-VAE, PepGlider w/o QN, PepGlider w/ *sign*, PepGlider w/ *z-norm*), described in Appendix A.4, and established VAE-based models for controlled AMP generation (HydrAMP (Szymczak et al., 2023), Transformer-128 (Renaud and Mansbach, 2023)), with detailed descriptions in Appendix A.5.

### 4.3 Metrics

We evaluate PepGlider’s performance across multiple dimensions during training and generation.

**Disentanglement Quality** is assessed using five established metrics: Interpretability, Correlation score, Modularity, Mutual Information Gap, and Separated Attribute Predictability (Pati and Lerch, 2021).

**Sequence Quality** is evaluated through: (1) *FBD*_*train*_, measuring conformity to training distribution via Fréchet Biological Distance with fine-tuned ESM2 embeddings as a validity proxy; (2) Diversity, quantified as the fraction of unique sequences not in the training set; (3) Novelty, computed via average pairwise Levenshtein distance; and (4) Fitness Score (Li et al., 2024), serving as a proxy for amphiphilicity, where higher scores indicate greater peptide functionality and antimicrobial activity.

**Biological Activity** assessment includes: (1) antimicrobial potency against *E. coli* and *S. aureus* using APEX predictions (Wan et al., 2024) for species-specific MIC values; (2) hemolytic toxicity via our trained classifier; and (3) *FBD*_*active*_, measuring similarity to experimentally validated antimicrobial peptides (MIC ≤ 32 *µ*g/ml).

Detailed evaluation methodology is provided in Appendix A.6.

## 5 Results

We evaluate PepGlider as continuous property regularization framework through studying fundamental controllability capabilities and domain-specific application. First, we assess core framework capabilities required for controllable peptide design, including **latent space disentanglement quality, continuous property control precision, and independent manipulation of correlated physicochemical properties**. Second, we demonstrate framework applicability through **antimicrobial peptide optimization**, showcasing how general controllability enables complex, domain-specific biological objectives. Generated peptides are evaluated using antimicrobial activity predictions, hemotoxicity assessment, and sequence quality metrics.

### 5.1 Disentanglement Quality

Effective disentanglement is crucial for controllable generation, as it determines whether individual properties can be manipulated independently through latent space traversal without unintended modifications of others at the same time. Following Pati and Lerch (2021), we assess PepGlider’s disentanglement quality using five established objective metrics: Interpretability, Correlation score (Corr score), Modularity, Mutual Information Gap (MIG), and Separated Attribute Predictability (SAP) averaged across charge, length, and hydrophobicity, presented in Table 1.

**Table 1:** Disentanglement quality metrics for PepGlider and baseline and ablation methods. Mean scores across peptide properties (charge, length, hydrophobicity). Higher scores indicate better disentanglement for all metrics. Property-specific results in Table S3.

| Model | Interpretability (↑) | Corr score (↑) | Modularity (↑) | MIG (↑) | SAP (↑) |
| --- | --- | --- | --- | --- | --- |
| VAE | 0.175 | 0.389 | 0.833 | 0.003 | 0.023 |
| HydrAMP | 0.231 | 0.487 | 0.864 | 0.012 | 0.025 |
| Transformer-128 | 0.104 | 0.365 | 0.845 | 0.005 | 0.039 |
| AR-VAE | 0.954 | **0.995** | 0.984 | 0.450 | 0.741 |
| PepGlider w/ signum | 0.955 | **0.995** | 0.984 | 0.453 | 0.739 |
| PepGlider w/o normalization | **0.981** | **0.995** | **0.987** | **0.479** | **0.771** |
| PepGlider w/ z-score normalization | 0.966 | **0.995** | **0.987** | 0.478 | 0.753 |
| PepGlider | **0.931** | **0.995** | **0.985** | **0.449** | **0.719** |

PepGlider achieves disentanglement quality comparable to the original AR-VAE across all metrics, demonstrating that our continuous loss formulation and quantile normalization preserve the strong disentanglement achieved through property-based regularization. All AR-VAE variants substantially outperform baseline methods (VAE, HydrAMP, Transformer-128), with 4 − 9× improvements in Interpretability (0.931 vs. 0.231) and 37-fold improvements in MIG (0.449 vs. 0.012). PepGlider maintains near-perfect monotonic alignment (Correlation score: 0.995) and performance comparable to AR-VAE across all metrics, with differences within 0.03 − 0.05. These results confirm that continuous property regularization with domain-specific normalization maintains latent space disentanglement, providing the foundation for quantitative property control.

#### 5.1.1 Continuous Property Control

Practical peptide design requires smooth, predictable property transitions during latent space traversal to enable systematic peptide optimization toward target properties values. We evaluate PepGlider’s continuous control capability by systematically sampling latent vectors, decoding them into sequences, and visualizing the resulting property values as 2D surfaces across pairs of latent dimensions.

The resulting property surfaces demonstrate smooth, continuous transitions across length, charge, and hydrophobicity (Figure 2, upper panel), enabling precise navigation through the latent space. The quality of latent space traversal is the highest for PepGlider when contrasted with baseline models (Figure S3). Simultaneously, PepGlider maintains consistently high validity, throughout latent space navigation, measured as FBD to training data (*FBD*_*train*_), shown in Figure 2 (lower panel). PepGlider’s consistent validity during traversal ensures that property optimization preserves biological plausibility of generated sequences. Additional amino acid frequency analysis confirms that PepGlider maintains realistic compositional patterns that closely match the training data distribution (Figure S4).

**Figure 2.**
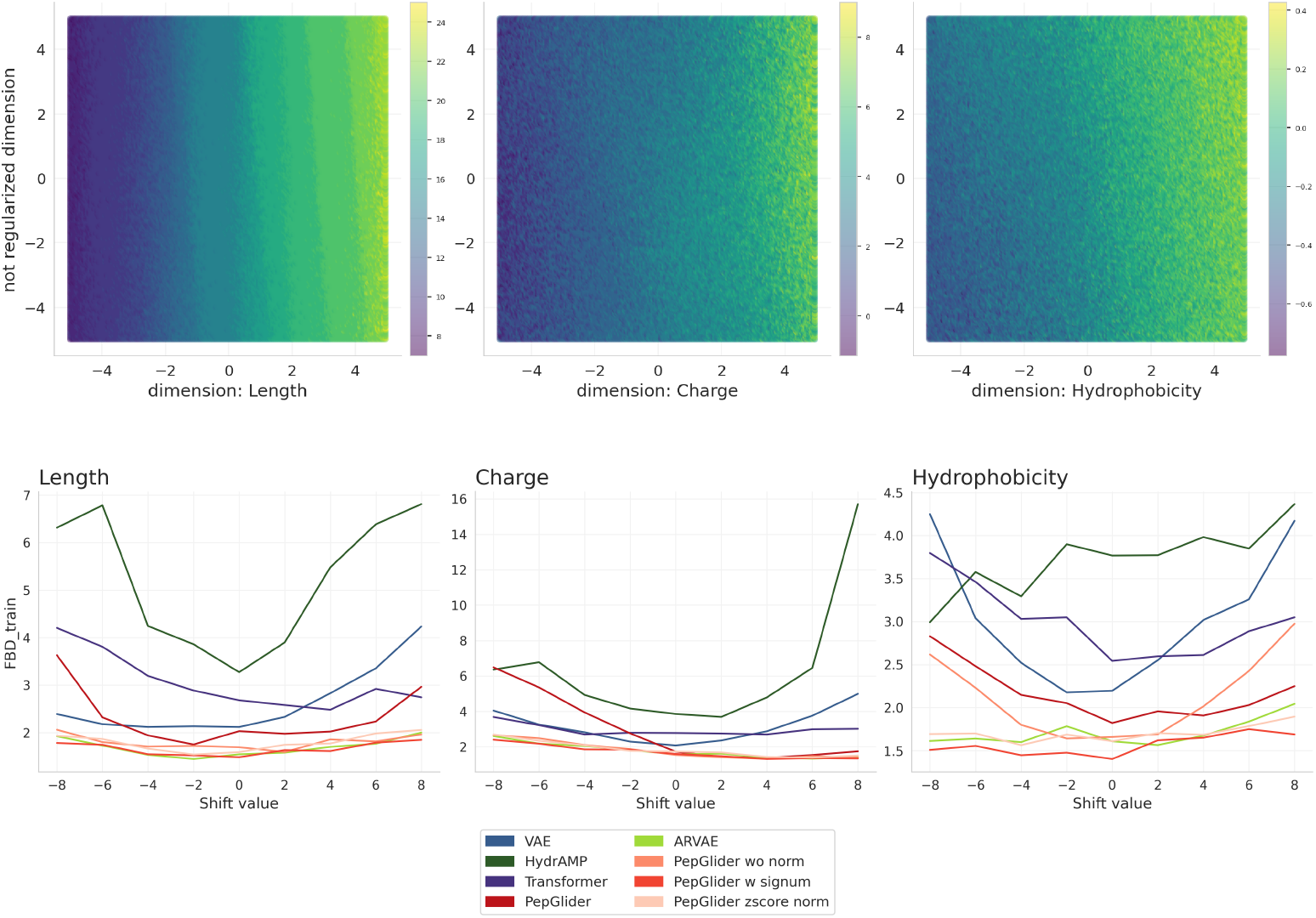
Continuous property control through latent space manipulation. **Upper panel:** PepGlider 2D property surface plots for length, charge, and hydrophobicity. **Lower panel:** Validity (*FBD*_*train*_) across latent shifts for PepGlider, baselines, and ablation variants.

#### 5.1.2 Independent Control of Correlated Properties

The ability to manipulate correlated properties addresses conflicting optimization objectives in peptide design. We evaluate PepGlider’s performance in this task through multi-property conditioning experiments, constraining different property combinations: individual physicochemicals properties (L, C, or H), pairs (L+C, L+H, or C+H), or all three simultaneously (L+C+H), measuring target property manipulation while monitoring cross-interference effects.

PepGlider achieves independent property control across multi-property conditioning scenarios (Figure 3). Single-property constraints provide linear control over the target property without affecting the non-target one. Multi-property conditioning maintains this selectivity, except for simultaneous charge and hydrophobicity control (C+H), where performance degrades due to physicochemical constraints, since hydrophobic residues are typically uncharged. Non-conflicting combinations (L+C, L+H) enable multi-objective control where each property responds to its corresponding latent dimension. Ablation analysis (Figure S5) shows PepGlider achieves the largest controllable value range across all variants, enabling more effective targeting of specific property values and exploration of otherwise inaccessible property regions.

**Figure 3.**
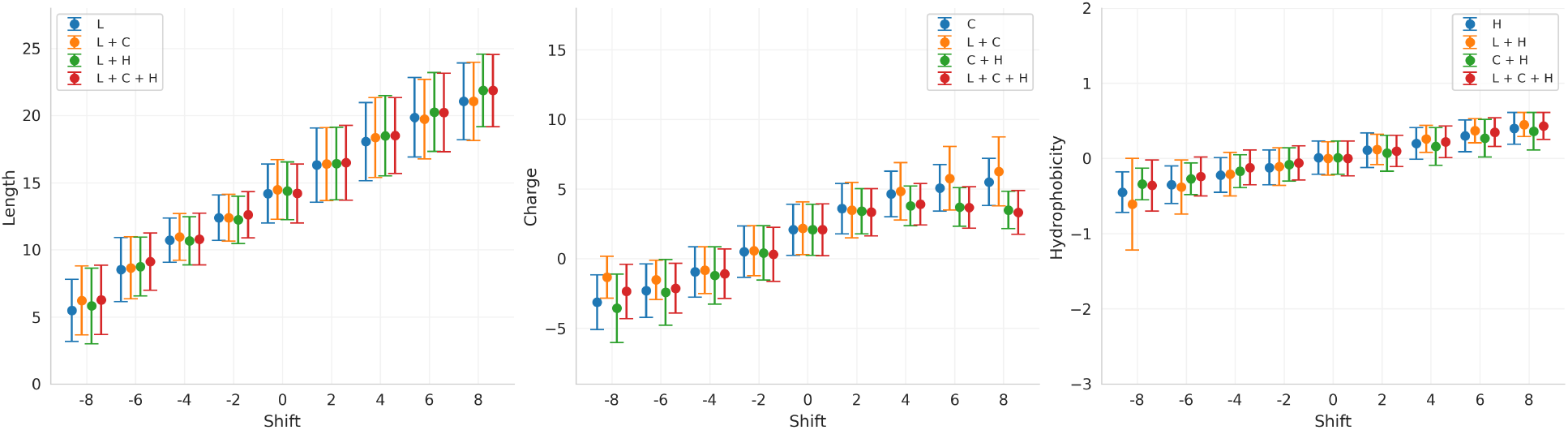
Independent control of correlated peptide properties through selective property conditioning. Property values across latent space displacements for **(a)** net charge (C), **(b)** sequence length (L), and **(c)** hydrophobicity (H) under single-property (blue), dual-property (orange, green), and tri-property (red) conditioning scenarios. Error bars represent mean ± standard deviation.

### 5.2 Antimicrobial Activity Optimization

To demonstrate practical applicability, we apply our framework to antimicrobial peptide optimization, where complex biological activity must be balanced against multiple physicochemical constraints. While for previously described experiments we used a model with regularized dimensions for basic peptide properties (length, charge, hydrophobicity), practical AMP design requires direct control over predicted biological activity We therefore evaluate two specialized models: one regularized for physicochemical properties (L, C, and H) and one for antimicrobial activity properties (predicted MIC against E. coli and S. aureus, non-toxicity). Analysis of generation quality, measured as *FBD*_*train*_ (Figure S6), reveals that specialization significantly improves performance within each model’s domain. At positive shift values, the antimicrobial-focused model achieves substantially lower *FBD*_*train*_ for MIC properties compared to the combined or physicochemical-only models. Similarly, the physicochemical-focused model shows superior stability for length, charge, and hydrophobicity throughout the controllable space. This separation enables us to apply the most appropriate model for each optimization objective: we use the antimicrobial-focused model for MIC optimization experiments, while physicochemical properties are controlled via the dedicated physicochemical model.

#### 5.2.1 Antimicrobial activity control

To evaluate whether PepGlider’s continuous control extends to complex biological functions, we generate 2D surface plots, where decoded peptides are evaluated using APEX MIC prediction models for *E. coli* and *S. aureus* or our toxicity classifier. The smooth activity gradients across latent space demonstrate systematic control over antimicrobial potency and hemotoxicity (Figure 4, upper panels). Validation through scatterplot analysis of in-house dataset peptides(described in Appendix A.2) projected into PepGlider’s latent space reveals that experimentally verified high-activity peptides (low MIC values) naturally cluster in regions associated with predicted antimicrobial efficacy (Figure 4, lower panels), confirming that learned representations capture genuine biological function rather than arbitrary encodings.

**Figure 4.**
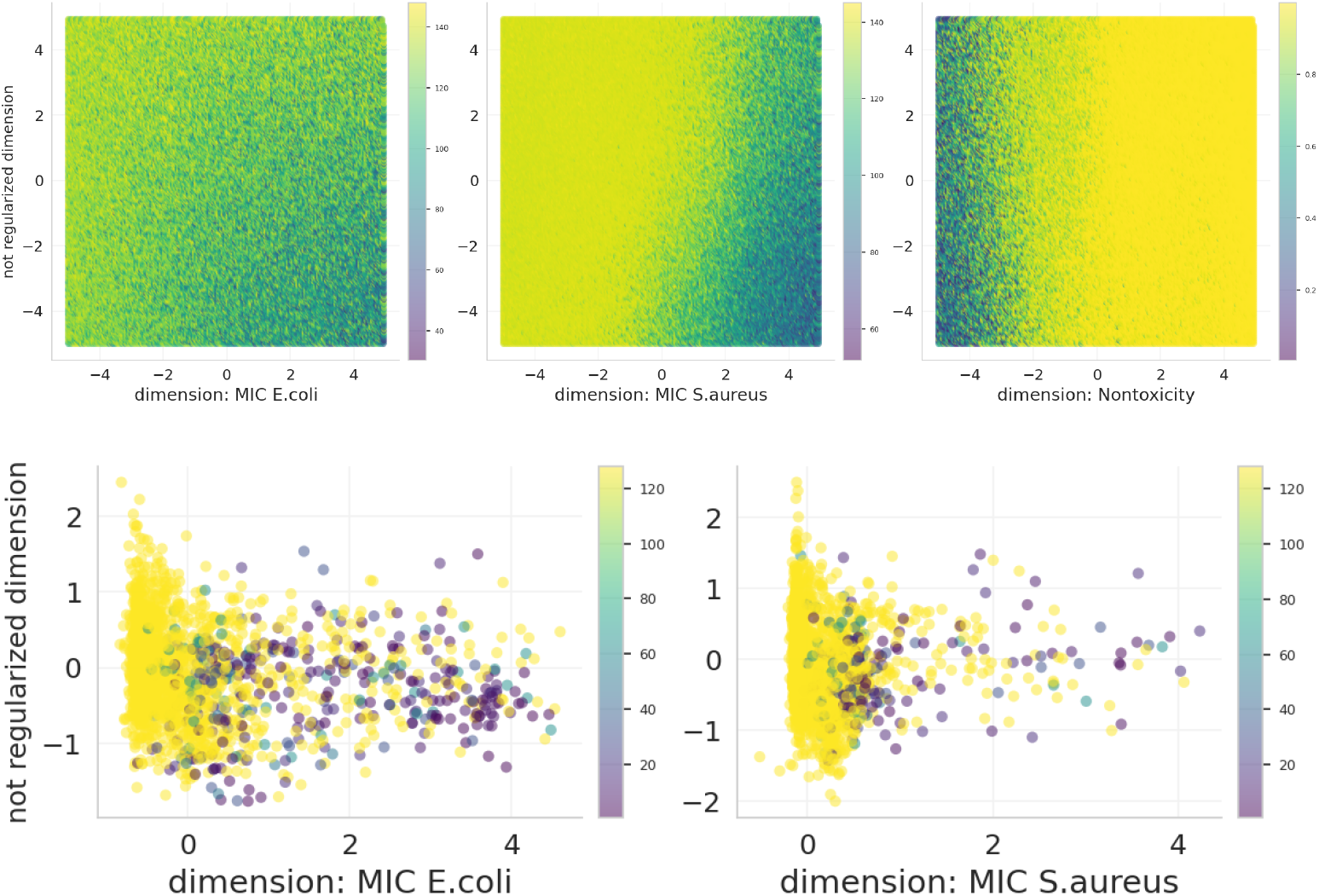
Continuous antimicrobial activity control in PepGlider latent space. **Upper panels:** 2D surface plots showing APEX-predicted MIC values across latent space for *E. coli* (left). *S. aureus* (middle), and non-toxicity predictions (right). **Lower panels:** Validation scatterplots showing peptides from proprietary dataset projected into latent space, colored by experimental MIC values for *E. coli* (left) and *S. aureus* (right).

#### 5.2.2 Unconstrained generation for high-activity peptide discovery

To evaluate PepGlider’s capability to discover high-activity peptides, we generate peptides in unconstrained mode (see Section 3.2.1) and assess their predicted activity and sequence quality. We compare against both our baseline models (VAE, HydrAMP, Transformer) and existing antimicrobial peptide generation methods from the literature: AMP-GAN (Van Oort et al., 2021), Diff-AMP (Wang et al., 2024), and AMP-Diffusion (Torres et al., 2025).

To identify optimal displacement values for latent dimensions regularized with respect to *E. coli* or *S. aureus* activity, we compare FBD to active peptides (*FBD*_*active*_) scores across different displacement values as a proxy for antimicrobial activity (Figure 5, upper panel). Based on these scores, we select optimal displacement values per strain (+8.0 for *E. coli*, +4.0 for *S. aureus*). At the same time, the validity of peptides generated for these values remains high (Figure S7). Using these strain-specific latent displacements, we generate sequences targeting each pathogen separately. Figure 5 (lower panel) shows APEX-predicted MIC distributions for these generated sequences, while Table 2 reports sequence quality metrics including *FBD*_*train*_ (validity), novelty, diversity, and fitness score.

**Table 2:**
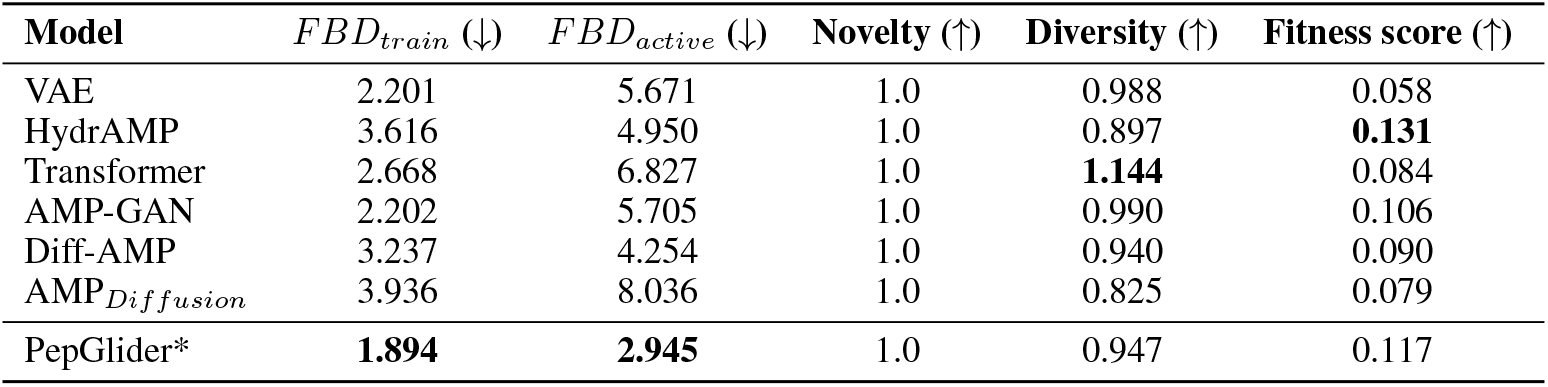
Sequence quality metrics for unconstrained generation across generative models. Comparison of *FBD*_*train*_, *FBD*_*active*_, novelty, and diversity for PepGlider and baseline/external generative methods (n=500). *PepGlider results averaged over *E. coli* and *S. aureus* predictions.

| Model | $FBD_{train}$ (↓) | $FBD_{active}$ (↓) | Novelty (↑) | Diversity (↑) | Fitness score (↑) |
| --- | --- | --- | --- | --- | --- |
| VAE | 2.201 | 5.671 | 1.0 | 0.988 | 0.058 |
| HydrAMP | 3.616 | 4.950 | 1.0 | 0.897 | **0.131** |
| Transformer | 2.668 | 6.827 | 1.0 | **1.144** | 0.084 |
| AMP-GAN | 2.202 | 5.705 | 1.0 | 0.990 | 0.106 |
| Diff-AMP | 3.237 | 4.254 | 1.0 | 0.940 | 0.090 |
| AMP<sub>Diffusion</sub> | 3.936 | 8.036 | 1.0 | 0.825 | 0.079 |
| PepGlider* | **1.894** | **2.945** | 1.0 | 0.947 | 0.117 |

**Figure 5.**
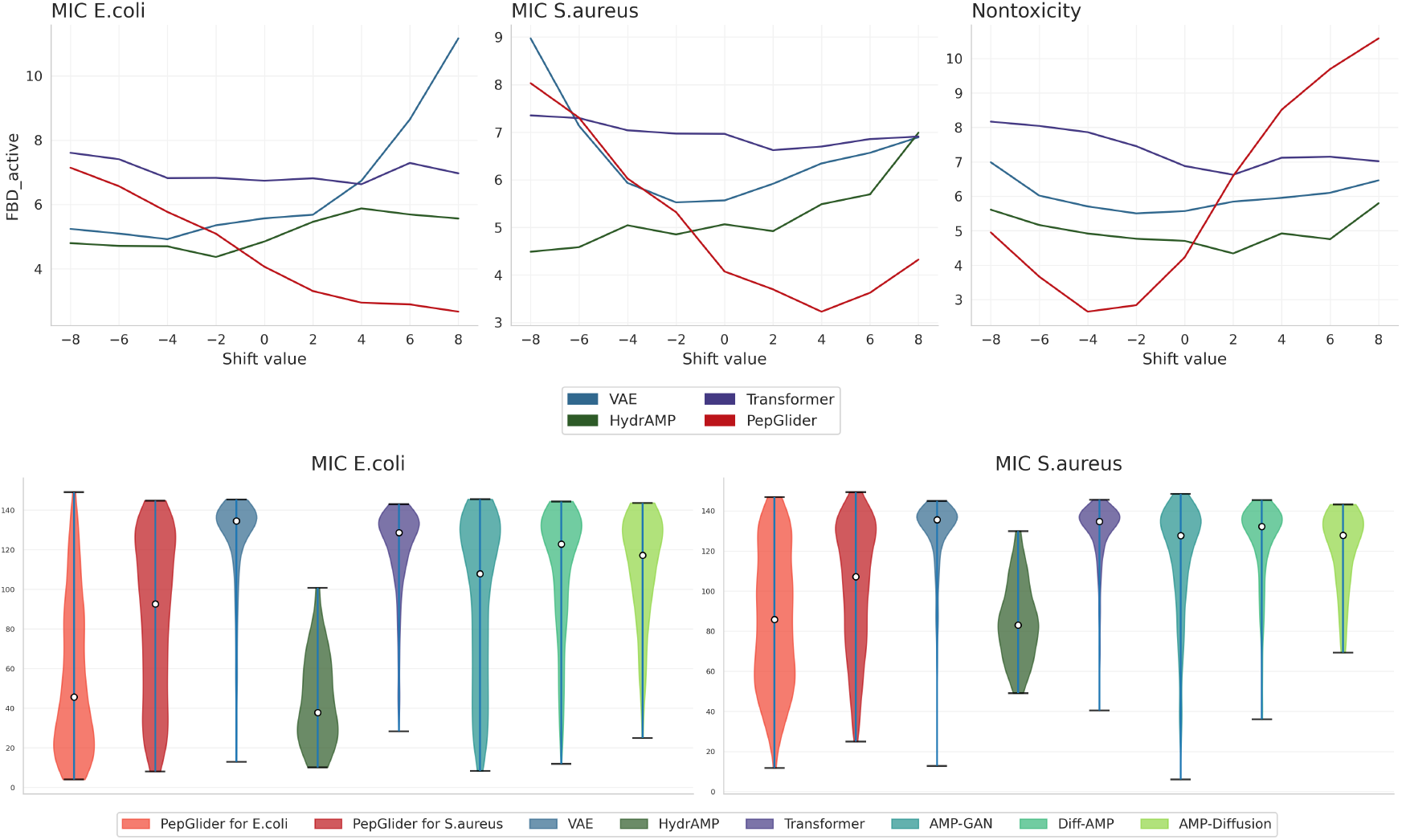
High-activity peptide generation through strategic latent space sampling. **(a)** *FBD*_*active*_ scores comparing PepGlider to baseline methods when generating peptides throughout latent space traversal. Lower *FBD*_*active*_ scores indicate greater similarity to genuine high-activity antimicrobial peptides. **(b)** MIC as predicted by APEX for *E. coli* (left) and *S. aureus* (right) from peptides generated by PepGlider when sampling from latent regions corresponding to low MIC predictions (*α* = +8 for *E. coli*, and *α* = +4 for *S. aureus*) and by baseline models (VAE, HydrAMP, Transformer, AMP-GAN, Diff-AMP, AMP-Diffusion).

PepGlider’s unconstrained generation produces antimicrobial candidates with substantially lower predicted MIC values than all comparison methods (Figure 5, lower panel). For *E. coli*, PepGlider generates sequences concentrated at 40 − 60 *µ*g/ml with significant density below 32 *µ*g/ml (clinically relevant threshold), while other methods show broader distributions centered at higher MIC values. For *S. aureus*, PepGlider maintains concentrated low MIC predictions.

HydrAMP demonstrates competitive performance among baseline methods, generating sequences with activity profiles between PepGlider and other models.

PepGlider achieves superior sequence quality across multiple metrics (Table 2). With *FBD*_*train*_ of 1.498, PepGlider generates sequences most similar to the training distribution while maintaining perfect novelty (no training set duplicates). PepGlider achieves the lowest *FBD*_*active*_ (2.935), indicating generated sequences most closely resemble genuine high-activity antimicrobial peptides. Diversity (0.956) remains comparable to baseline methods, demonstrating that PepGlider’s enhanced activity targeting does not compromise sequence variation. Notably, PepGlider achieves competitive fitness scores (0.117), ranking second among all methods and indicating strong amphiphilic character essential for membrane disruption. This emergent property, though not explicitly controlled during training, arises naturally from regularization of correlated antimicrobial activity. These results confirm that continuous property regularization enables effective biological activity enhancement through unconstrained generation.

#### 5.2.3 Analog generation for optimizing AMP activity

We evaluate PepGlider’s analog generation through targeted manipulation of regularized latent dimensions, comparing against HydrAMP, the only baseline model supporting analog generation. Starting from prototype sequences classified as active (MIC ≤ 32 *µ*g/ml) or inactive (MIC ≥ 128 *µ*g/ml), we modify existing peptides to enhance predicted activity against *E. coli* or *S. aureus*. PepGlider applies displacement values (*α* = 1, 4, 8) to regularized dimensions, exploiting the monotonic relationship between latent dimension values and predicted MIC to control activity enhancement. HydrAMP uses temperature parameters (*τ* = 1, 2.5, 5) with conditioning toward high activity.

PepGlider demonstrates superior activity enhancement across both bacterial targets (Figure 6). Starting from both active and inactive prototypes, PepGlider consistently achieves lower predicted MIC values than HydrAMP across all displacement values, with *α* = 8 reaching clinically relevant potency around 32 *µ*g/mL. PepGlider shows robust improvement even from inactive peptides, effectively transforming low activity sequences into promising candidates. *S. aureus* activity improvement proves more challenging than *E. coli*, particularly from already active prototypes. PepGlider enables independent optimization for each bacterial strain through separate regularized dimensions, allowing strain-specific targeting within a single framework. This capability is rarely achieved in antimicrobial peptide design where multi-strain optimization typically requires distinct models or *post-hoc* filtering approaches (Szymczak and Szczurek, 2023).

**Figure 6.**
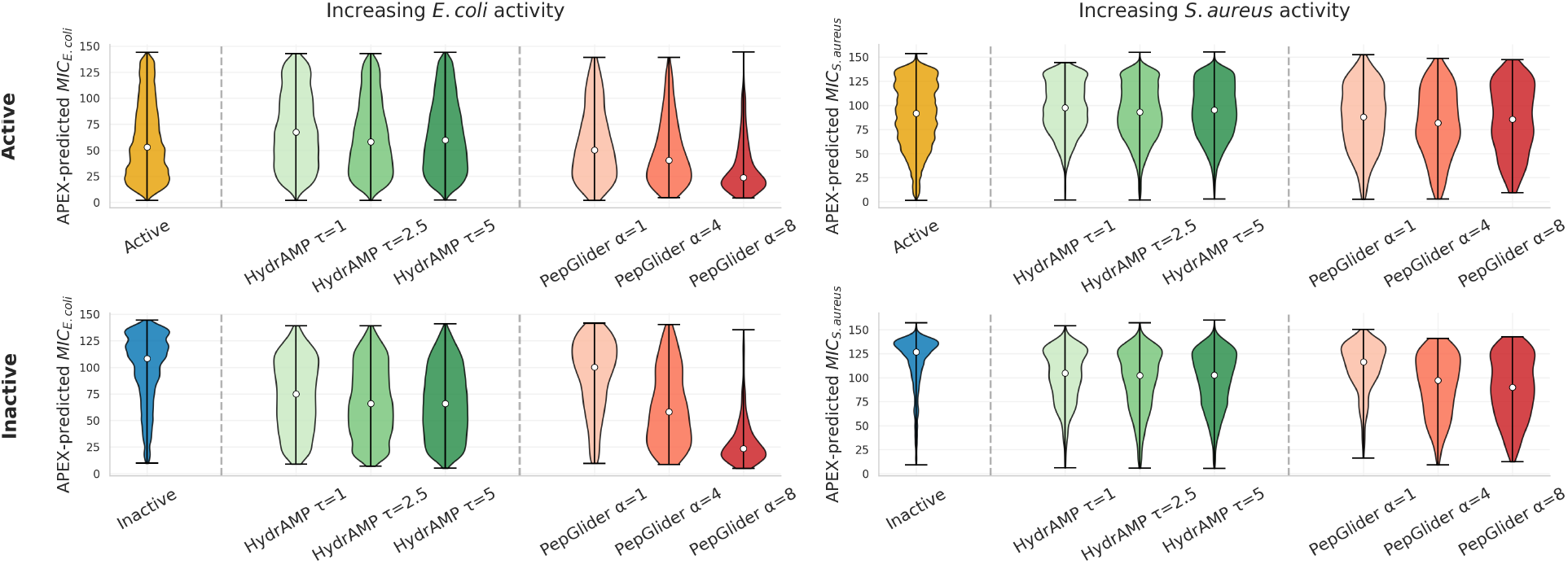
Peptide activity optimization through analog generation. Violin plots showing APEX-predicted MIC distributions for PepGlider and HydrAMP analog generation across active prototypes (MIC ≤ 32 *µ*g/ml, upper panel) and inactive prototypes (MIC ≥ 128 *µ*g/ml, lower panel) for *E. coli* (left) and *S. aureus* (right). HydrAMP parameters: temperature *τ* ∈ {1, 2.5, 5}. PepGlider parameters: displacement values *α* ∈ {1, 4, 8} applied to regularized dimensions corresponding to strain-specific activity.

#### 5.2.4 Multi-objective optimization of antimicrobial activity and hemotoxicity

A critical challenge in antimicrobial peptide development is balancing efficacy against safety, as enhanced activity often correlates with increased toxicity. We evaluate PepGlider’s ability to navigate this trade-off through simultaneous manipulation of MIC and non-toxicity regularized dimensions. While applying positive displacements to MIC-regularized dimensions to enhance antimicrobial activity against *E. coli* and *S. aureus*, we fix the non-toxicity regularized dimension at specific values 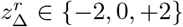 to assess whether toxicity increases can be mitigated during activity enhancement.

PepGlider demonstrates capacity for controlled multi-objective optimization through simultaneous manipulation of activity and toxicity dimensions. As positive displacements are applied to MIC-regularized dimensions (shift values 0 to +8), predicted antimicrobial activity increases for both *E. coli* and *S. aureus*, with MIC values decreasing from approximately 140 *µ*g/ml to below 60 *µ*g/ml (Figure 7, left panels). The non-toxicity dimension setting directly controls the activity-safety trade-off (Figure 7, right panels). When the non-toxicity dimension is fixed at 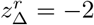, sequences achieve the lowest MIC values, reaching approximately 30 *µ*g/ml at high shift values, but this comes at the cost of substantial non-toxicity degradation, with scores dropping below 0.7. Conversely, fixing the non-toxicity dimension at 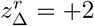 maintains non-toxicity scores above 0.95 across all activity levels, though with slightly higher MIC values.

**Figure 7.**
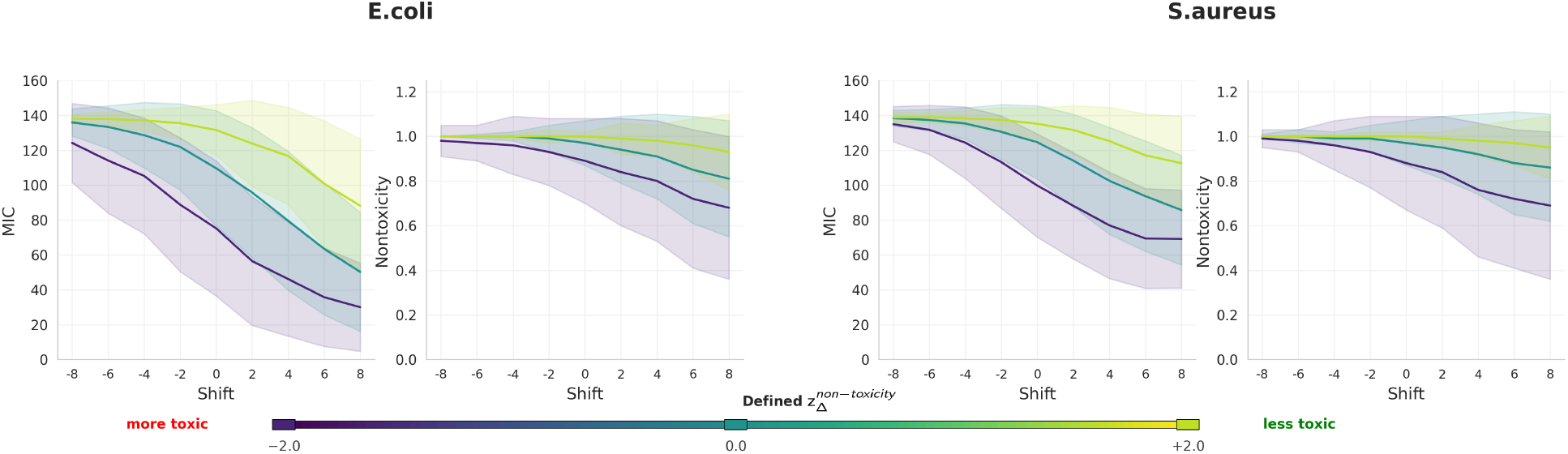
Multi-objective optimization of antimicrobial activity and non-toxicity. MIC predictions for *E. coli* (upper left) and *S. aureus* (lower left) when manipulating respective MIC-regularized dimensions, with simultaneous non-toxicity predictions under different non-toxicity regularization strategies: *α* = −2 (violet), *α* = 0 (blue), *α* = +2 (yellow). Mean trajectories (solid lines) with ±1 standard deviation confidence bands (shaded regions) for predicted antimicrobial activity and non-toxicity under varying distribution shifts. Color mapping follows the 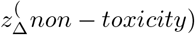 parameter space, where purple (*z* =− 2.0) represents toxic shifts and yellow-green (*z* = +2.0) represents non-toxic shifts.

The neutral setting 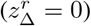 shows intermediate behavior, with gradual non-toxicity decline as activity increases. These results confirm that PepGlider enables independent manipulation of correlated properties, allowing enhancement of antimicrobial activity while explicitly controlling toxicity through separate regularized dimensions.

## 6 Discussion

PepGlider addresses fundamental limitations in controllable peptide design through continuous attribute regularization and adaptive normalization strategies. Our framework demonstrated superior latent quality evaluation and systematic property control across challenging scenarios like activity-safety trade-offs.

Key limitations include reduced performance for inherently conflicting objectives, particularly the activity-toxicity trade-off where enhanced antimicrobial potency often correlates with increased toxicity. Future developments should prioritize methods that efficiently utilize sparse biological experimental data directly, reducing dependence on intermediate prediction models while maintaining controllability. The current set, while comprehensive for basic physicochemical properties, could be expanded to include synthesizability constraints, structural features (secondary structure propensity, flexibility), and manufacturing considerations critical for therapeutic translation.

The continuous property regularization framework’s versatility extends beyond antimicrobial peptides to diverse therapeutic applications, providing a flexible framework for controllable generation. including self-assembling peptides for materials science or cell-penetrating peptides for drug delivery, establishing a generalizable methodology for precise property control across the broader peptide design landscape.

### Ethics statement

This research involves computational design of antimicrobial peptides using machine learning methods. All datasets used for training and evaluation consist of publicly available peptide sequences and experimental measurements from established databases. No human subjects, animal experiments, or clinical trials were involved in this computational study. The potential therapeutic applications of designed antimicrobial peptides could contribute to addressing antimicrobial resistance, a significant global health challenge. However, any peptides generated by this framework require extensive experimental validation, safety testing, and regulatory approval before clinical consideration. The hemolytic toxicity predictions used in this work are computational estimates and cannot replace experimental safety assessment.

### Reproducibility statement

We provide comprehensive implementation details to ensure reproducibility. Model architectures, hyperparameters, and training procedures are detailed in the main text and appendix (Table S1). All normalization procedures, including quantile transformation and adaptive range normalization, are mathematically specified with explicit equations. Evaluation metrics and baseline comparisons use established methods with clear mathematical definitions. The proprietary validation dataset contains experimental MIC measurements that enable independent performance assessment, though specific data cannot be shared due to proprietary restrictions. Code and trained models will be made available upon publication to facilitate reproduction and extension of this work.

### LLM usage

Large language models were used as writing assistance tools during the preparation of this manuscript. Specifically, Claude was employed for improving clarity and flow of technical explanations, and generating alternative phrasings for complex methodological concepts, proofreading and copy-editing assistance. All scientific content, experimental design, results, and conclusions are entirely the work of the human authors. LLMs were not used for experimental design decisions, or scientific reasoning. The core contributions, methodological innovations, and technical implementations represent original research by the authors. All factual claims and experimental results were verified independently by the research team.

## Acknowledgments

The authors gratefully acknowledge Kamil Deja, Tomasz Trzciński and the Technical University of Warsaw for generously providing access to computational resources that enabled this work. This research was supported in part by PL-Grid Infrastructure grant nr PLG/2025/018424.

## Author Contributions

**Aleksandra Gwiazda**: Conceptualization, Software, Investigation, Validation, Visualization, Writing - Review & Editing

**Paulina Szymczak**: Conceptualization, Methodology, Formal analysis, Data curation, Visualization, Writing - Original Draft, Project administration

**Ewa Szczurek**: Writing - Review & Editing, Supervision, Funding acquisition

## A Appendix

### A.1 Extended Experimental Setup

#### A.1.1 Physicochemical properties

We selected net charge, hydrophobicity, and sequence length as target properties for PepGlider based on their established roles in antimicrobial peptide function and their ability to discriminate between active and inactive peptides.

- **Net Charge** plays a critical role in the initial electrostatic interactions between cationic antimicrobial peptides and negatively charged bacterial membranes (Yeaman and Yount, 2003). Positively charged residues facilitate binding to bacterial lipopolysaccharides and phospholipids, while excessive charge can lead to reduced membrane permeation and potential cytotoxicity (Hancock and Sahl, 2006). Net charge is calculated at physiological pH (7.0), accounting for the ionization state of terminal groups and ionizable side chains based on their respective pKa values.
- **Hydrophobicity** determines the peptide’s ability to partition into and disrupt bacterial membranes (Wieprecht et al., 1997). Optimal hydrophobic content enables membrane insertion while preventing aggregation or excessive hemolytic activity. Average hydrophobicity is computed using established amino acid hydrophobicity scales:

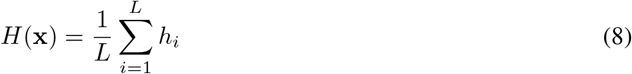

where *h*_*i*_ represents the hydrophobicity value for amino acid *i* and *L* is the sequence length.
- **Sequence length** constrains both structural flexibility and membrane interaction mechanisms. Shorter peptides typically adopt extended conformations that facilitate membrane carpet formation, while longer sequences may form more complex secondary structures affecting activity and selectivity (Shai and Oren, 2001). Sequence length is a direct enumeration of amino acid residues:

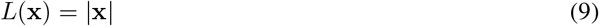

Physicochemical properties are computed using the *modlamp* (Müller et al., 2017) *package, implementing algorithms for antimicrobial peptide analysis*.

To validate these properties as discriminative features, we analyzed their distributions across **active** (MIC ≤ 32 *µ*g/ml) and **inactive** (MIC ≥ 128 *µ*g/ml) peptides in our curated dataset (Figure S1). Active peptides exhibit distinct distributions for all three properties: moderate positive charge, intermediate hydrophobicity values, and concentrated length distributions around 10-25 residues. These clear distributional differences support their selection as target properties for continuous control in the PepGlider framework.

#### A.1.2 Antimicrobial activity

##### Minimum Inhibitory Concentration (MIC)

MIC represents the lowest concentration of an antimicrobial agent that prevents visible bacterial growth after a defined incubation period, serving as the gold standard for quantifying antimicrobial potency. MIC values are expressed in *µ*g/ml, with lower values indicating higher antimicrobial activity. Clinical breakpoints typically classify peptides with MIC ≤ 32 *µ*g/ml as having therapeutically relevant activity (Hancock and Sahl, 2006), while values ≥ 128 *µ*g/ml indicate minimal to no antimicrobial effect.

For dataset analysis and validation purposes, we define **active peptides** as those achieving MIC ≤ 32 *µ*g/ml against at least one bacterial strain, indicating therapeutically relevant antimicrobial activity. Conversely, **inactive peptides** are defined as those with MIC ≥ 128 *µ*g/ml against all tested strains, indicating insufficient antimicrobial potency. Experimental MIC measurements for active/inactive classification and model validation are obtained from the DBAASP (Database of Antimicrobial Activity and Structure of Peptides) database (Pirtskhalava et al., 2021).

**Supplementary Figure S1:**
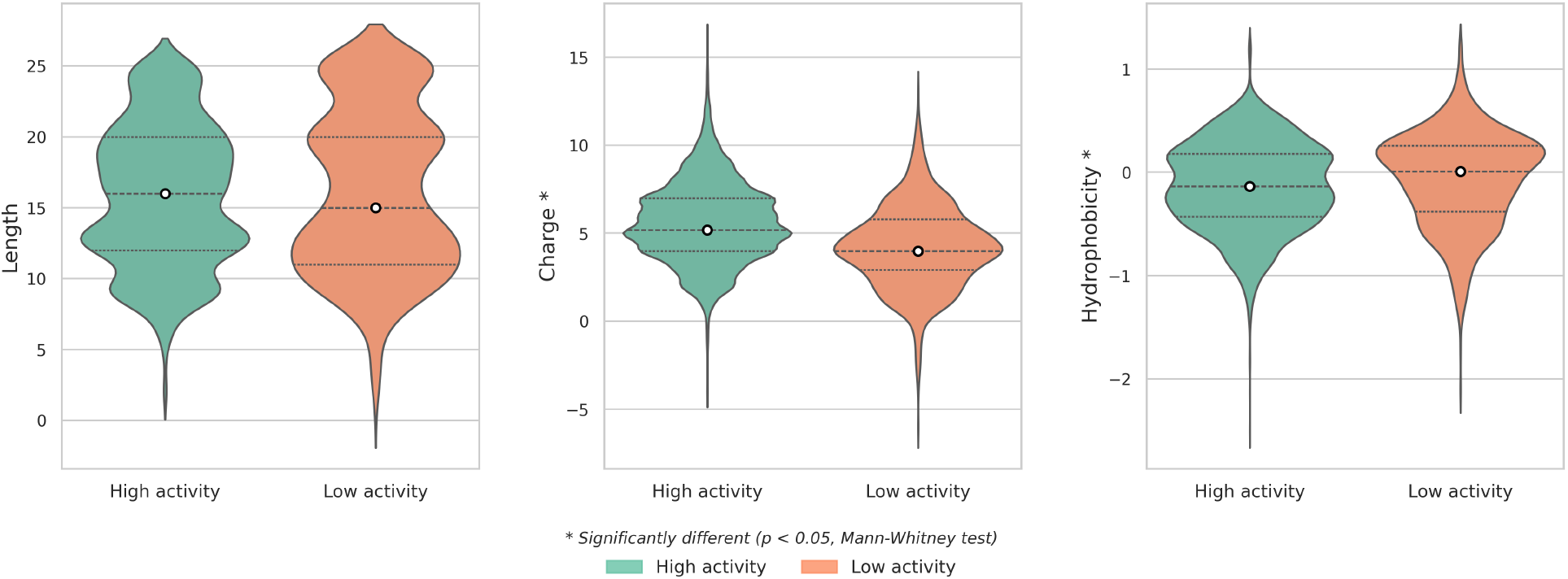
Discriminative physicochemical properties between active and inactive antimicrobial peptides. Distribution comparison of net charge, hydrophobicity, and sequence length between active peptides (green) and inactive peptides (orange) from the curated dataset.

##### APEX Antimicrobial Activity Predictor

We employ APEX (Wan et al., 2024), a deep learning framework trained on experimentally validated MIC measurements from over 3,000 antimicrobial peptides tested against 11 clinically relevant bacterial strains. APEX employs an encoder neural network combining recurrent and attention mechanisms for peptide feature extraction, coupled with strain-specific downstream regression networks for MIC prediction. APEX predictions serve as continuous MIC values for PepGlider regularization, allowing direct optimization of antimicrobial potency through latent space manipulation.

##### Target Bacterial Strains

We focus on two clinically relevant bacterial pathogens representing distinct membrane architectures and infection profiles:

- ***Escherichia coli* (*E. coli*)**: A Gram-negative bacterium with an outer membrane containing lipopolysaccharides, representing a major cause of urinary tract infections, sepsis, and antibiotic-resistant nosocomial infections. We average MIC predictions across three strains: ATCC 11775 (reference strain), AIG222 (carbapenem-resistant), and AIG221 (multidrug-resistant). This averaging captures activity against both susceptible and resistant *E. coli* variants.
- ***Staphylococcus aureus* (*S. aureus*)**: A Gram-positive pathogen with a thick peptidoglycan cell wall, responsible for skin infections, pneumonia, and life-threatening methicillin-resistant *S. aureus* (MRSA) infections. We average predictions across ATCC 12600 (reference strain) and ATCC BAA-1556 (MRSA strain), ensuring generated peptides target both susceptible and clinically critical resistant variants.

Strain-averaged MIC values provide robust activity estimates while reducing sensitivity to strain-specific variations, enabling PepGlider to optimize for broad-spectrum antimicrobial activity within each bacterial species.

#### A.1.3 Non-toxicity

##### Data Extraction and Preprocessing

Hemolytic activity measurements were extracted from DBAASP (Pirtskhalava et al., 2021), focusing on records containing HC50 values and percentage hemolysis data. To increase the size of the non-toxic dataset, we included metabolic (n=36882), signal (n=868336), and hormone peptides (n=65087) extracted from Peptipedia Cabas-Mora et al. (2024) (total of 970305) peptides, which are presumed non-hemolytic due to their physiological roles. Activity measure values and groups were normalized to lowercase for consistent processing. Percentage hemolysis values were extracted using regex parsing with two prioritized patterns: (1) values with standard deviations (e.g., “15.2±3.1% hemolysis”), taking the primary value before ±, and (2) range formats (e.g., “10-20% hemolysis”), using the midpoint average. The resulting dataset contains 1157 hemolytic and 3120 non-hemolytic sequences.

##### Binary Toxicity Classification Rules

For peptides with single measurements, the following hierarchy was applied:

- Direct non-toxic assignment: 0% hemolysis records → non-toxic (1)
- Activity-based toxic assignment: activity ≤32 *µ*g/mL AND *>*1% hemolysis → toxic (0)
- Activity-based non-toxic assignment: activity *>*32 *µ*g/mL AND ≤10% hemolysis → non-toxic (1)
- Hemolysis threshold: *>*50% hemolysis → toxic (0)
- HC50-specific rules: HC50 ≤256 *µ*g/mL → toxic (0), otherwise non-toxic (1)

For peptides with multiple measurements, consensus rules were applied:

- Any measurement *<*32 *µ*g/mL → toxic (0)
- All measurements *>*128 *µ*g/mL → non-toxic (1)
- All measurements ≤10% hemolysis → evaluate based on activity (*<*32 *µ*g/mL threshold for toxic (0))
- HC50 measurements prioritized when available, using 128 *µ*g/mL threshold (toxic (0) if ≤128 *µ*g/mL)

Peptides not meeting any classification criteria were excluded from the training dataset.

##### Feature Engineering

The physicochemical property calculation framework computed 100+ features per sequence, including:

- Basic properties: length, charge, isoelectric point, molecular weight, aromaticity
- Hydrophobicity scales: AASI, Argos, Eisenberg, GRAVY, Kyte-Doolittle (16 scales total)
- Structural descriptors: secondary structure fractions, flexibility, entropy
- Specialized scales: Z-scales (5D), Kidera factors (10D), VHSE scales (8D), FASGAI vectors (6D)
- Topological features: polar surface area, H-bond acceptors/donors, rotatable bonds
- Compositional features: amino acid frequencies, structural class distributions

##### Model Training and Validation

XGBoost classifier hyperparameters were optimized on the training set with 974,582 peptides (1,157 toxic and 973,425 non-toxic) balanced using focal loss set to handle the imbalanced dataset. Model performance was assessed on the independent HydrAMP dataset containing experimentally validated antimicrobial peptides with known hemolytic profiles, and obtained accuracy of 0.83, and F1-Score of 0.90).

#### A.1.4 Property Normalization Procedures

We introduce property-specific normalization strategies as a core methodological contribution that ensures compatibility with our continuous loss formulation while preserving biological meaning.

##### Quantile Normalization

Applied to charge, length, hydrophobicity, and non-toxicity predictions. Raw values are transformed via quantile transformation *Q*(*·*) to uniform distribution *U* (0, 1), then linearly scaled:

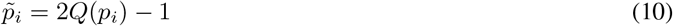

This ensures uniform property space coverage and eliminates scale bias while maintaining the required [−1, 1] range.

##### Adaptive Range Normalization

A normalization strategy that addresses the clinical importance of low MIC values while maintaining loss compatibility. The approach allocates 70% of the normalized range to therapeutically relevant concentrations (0-32 *µ*g/ml) and 30% to higher values. This 70/30 split was chosen empirically to balance sufficient resolution for precise control within the therapeutic range while maintaining representation of inactive peptides:

*Higher concentrations (>32 µg/ml)* → *[−1, −0*.*4]:*

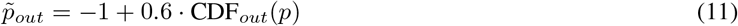

*Clinically relevant range (0-32 µg/ml)* → *[-0*.*4, 1]:*

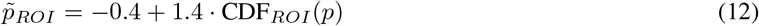

where CDF_*ROI*_ and CDF_*out*_ are empirical cumulative distribution functions computed within each region using histogram-based quantile mapping.

This normalization framework enables precise property control by ensuring all normalized properties operate within the same bounded range as the latent regularization terms.

**Supplementary Figure S2:**
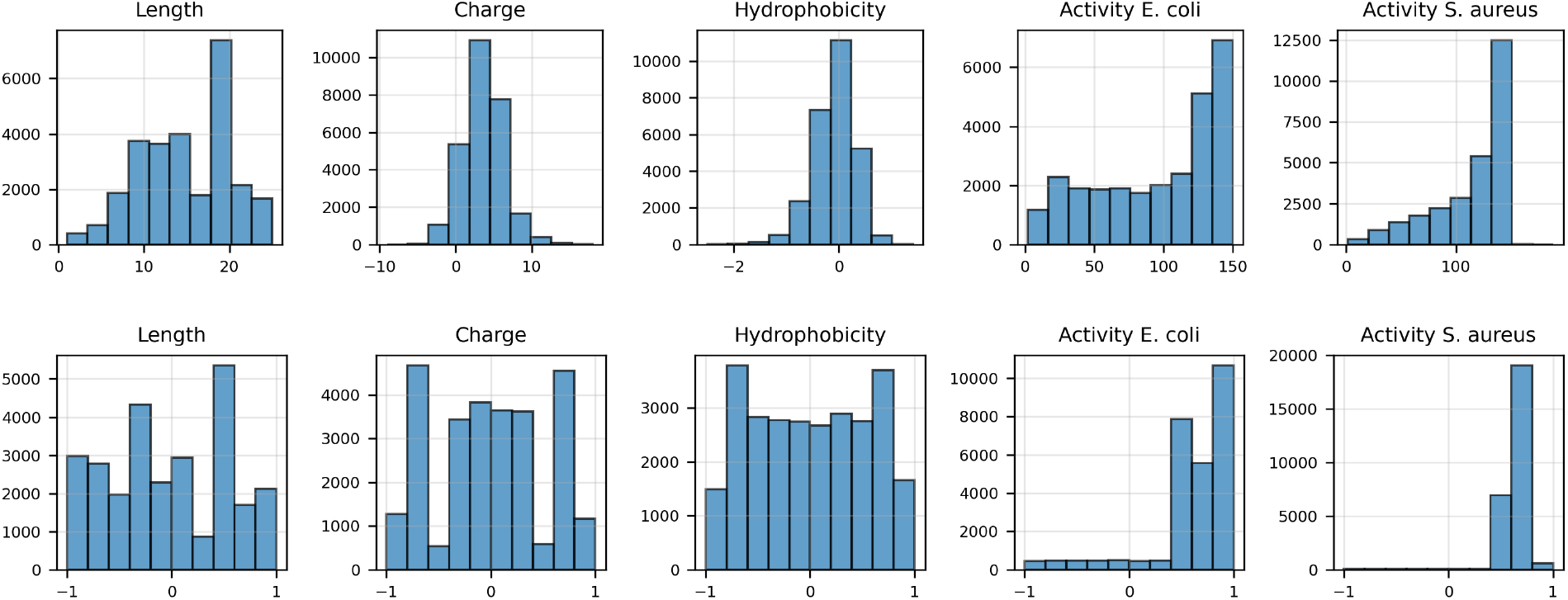
Normalization procedure for continuous property regularization. Distribution comparison of peptide properties before (upper panel) and after (lower panel) the normalization procedure.

### A.2 Datasets

#### PepGlider Training Dataset

The training dataset comprises 27331 curated antimicrobial peptide sequences derived from four comprehensive databases: AMPScanner (Veltri et al., 2018), dbAMP (Yao et al., 2025), DRAMP (Kang et al., 2019), and DBAASP (Pirtskhalava et al., 2021). Sequences are restricted to a maximum of 25 amino acid residues, capturing the predominant length range of naturally occurring AMPs. Duplicate sequences are removed across databases to ensure unique representation within the training corpus.

#### Evaluation Dataset

For evaluation of PepGlider, we use a proprietary dataset containing experimental MIC measurements for 1,736 peptides tested against 11 clinically relevant bacterial strains. The dataset includes measurements against Gram-negative bacteria (*Acinetobacter baumannii* (*A. baumannii*) ATCC 19606, *E. coli* ATCC 11775, *E. coli* AIC221, carbapenem-resistant *E. coli* AIC222, *Klebsiella pneumoniae* (*K. pneumoniae*) ATCC 13883, *Pseudomonas aeruginosa* (*P. aeruginosa*) PAO1 and PA14) and Gram-positive bacteria (*S. aureus* ATCC 12600, MRSA ATCC BAA-1556, vancomycin-resistant *Enterococcus faecalis* (*E. faecalis*) ATCC 700802, and vancomycin-resistant *Enterococcus faecium* (*E. faecium*) ATCC 700221).

### A.3 PepGlider implementation

PepGlider employs a transformer-based VAE architecture (Kingma and Welling, 2013) optimized for variable-length biological sequences. The encoder Enc(*·*) maps peptide sequences **x** ∈ {*A, C, D*, …, *Y*} ^*L*^ to latent representations **z** ∈ ℝ^*d*^ through CLS token aggregation, where a learnable classification token attends to all sequence positions via multi-head self-attention mechanisms. The encoder outputs parameterize a Gaussian posterior *q*(**z** | **x**) with mean ***µ***(**x**) and standard deviation ***σ***(**x**). The decoder Dec(*·*) reconstructs sequences from latent codes by replicating the latent vector across sequence positions and applying positional encodings for position-specific token generation.

The architecture incorporates *β*-VAE regularization (Higgins et al., 2017) and Importance Weighted Autoencoder components (Burda et al., 2015) to enhance posterior distribution approximation and latent space disentanglement capabilities. The *β* and *γ* parameter follow linear annealing schedules from their initial to final values over the specified warmup steps, after which they remain constant. The KL divergence weight *β* gradually increases to prevent posterior collapse.

All models were trained on NVIDIA A100 GPUs with 8GB memory. PepGlider and baseline models were trained for approximately 48 hours. The hyperparameter details are presented in Table S1.

**Supplementary Table S1:** Model hyperparameters and training details.

| Parameter | Value |
| --- | --- |
| **Architecture** |  |
| Attention Heads | 4 |
| Transformer Layers | 6 |
| Latent Dimension | 56 |
| Positional Encoding | Additive |
| Dropout Rate | 0.1 |
| Layer Normalization | Enabled |
| **Training** |  |
| Optimizer | Adam |
| Learning Rate | 0.001 |
| Batch Size | 512 |
| Epochs | 3100 |
| IWAE Samples ($K$) | 10 |
| **VAE Regularization** |  |
| $\beta$ Initial | 0.00001 |
| $\beta$ Final | 0.1 |
| $\beta$ Warmup Steps | 8000 |
| **Property Regularization** |  |
| Regularized Dimensions | [0, 1, 2, 3, 4, 5] |
| $\gamma$ Initial | 0.00001 |
| $\gamma$ Final | 20 |
| $\gamma$ Warmup Steps | 8000 |
| $\gamma$ Triggered epoch | 1000 |
| $\delta$ | 0.1 for PepGlider for physicochemical properties<br>0.6 for PepGlider only antimicrobial properties |
| **Data** |  |
| Max Sequence Length | 25 |
| Vocabulary Size | 20 |
| Property Normalization | Quantile (10 bins) |
| Properties Regularized | Length, Charge, Hydrophobicity, MIC E.coli, MIC S.aureus, Non-toxicity |

### A.4 Ablations

**VAE Baseline** serves as our primary ablation control, employing the identical transformer-based VAE architecture as PepGlider with Importance Weighted Autoencoder components and *β*-VAE regularization. The model is trained on the same dataset with identical quantile normalization, but without the continuous property regularization loss (*γ* = 0) and 10-times decreased *β*. This configuration isolates the contribution of our continuous regularization framework from the architectural and preprocessing components.

**AR-VAE** (Pati and Lerch, 2021) represents the original attribute regularization formulation using the signum function for discrete ordinal comparisons and standard preprocessing without quantile normalization. We implement AR-VAE using our transformer-based encoder and decoder architecture for fair comparison. This baseline evaluates the impact of our continuous loss formulation and normalization improvements while controlling for architectural differences.

**PepGlider w/o QT** removes quantile normalization while retaining the continuous loss formulation and signum replacement. Raw property values are used directly in the loss computation, isolating the contribution of the normalization procedure.

**PepGlider w/ sign** retains the original signum function from AR-VAE while incorporating our quantile normalization approach. This variant evaluates whether normalization alone can improve discrete property regularization.

**PepGlider w/ z-norm** replaces quantile normalization with standard z-score normalization (zero mean, unit variance), testing alternative normalization strategies while maintaining the continuous loss formulation.

### A.5 Baseline Models

**HydrAMP** (Szymczak et al., 2023) is a conditional variational autoencoder for antimicrobial peptide generation. The model employs Jacobian-based disentanglement regularization to enforce independence between latent representations z and discrete conditioning variables (*c*_*AMP*_, *c*_*MIC*_ ∈ { 0, 1}). Property control is achieved through conditional decoding Dec(**z**, *c*) with binary labels for antimicrobial activity and potency. In contrast to PepGlider’s continuous property regularization in latent space, HydrAMP guides generation through discrete conditions fed directly to the decoder. The model supports unconstrained generation and temperature-controlled analogue generation modes.

**Transformer-128**

(Renaud and Mansbach, 2023) employs a transformer-based autoencoder architecture with a 128-dimensional latent space for peptide generation. The model learns implicit partitioning of the latent space into regions corresponding to high and low AMP probabilities without explicit incorporation of mechanisms for continuous property control.

### A.6 Evaluation

#### A.6.1 Disentanglement Quality

Following Pati and Lerch (2021), we assess PepGlider’s disentanglement quality using five established objective metrics:

- **Interpretability** measures how well individual latent dimensions align with specific properties by evaluating the variance explained by the most informative dimension for each attribute:

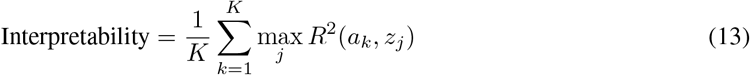

where *K* is the number of attributes, *a*_*k*_ is the *k*-th attribute, and *R*^2^(*a*_*k*_, *z*_*j*_) is the coefficient of determination between attribute *k* and latent dimension *j*.
- **Correlation score (Corr score)** quantifies the monotonic relationship between latent dimensions and target attributes by measuring the strongest rank correlation between each attribute and any latent dimension using Spearman’s correlation coefficient:

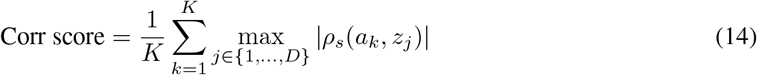

where *K* is the number of attributes, *D* is the number of latent dimensions, *ρ*_*s*_(*a*_*k*_, *z*_*j*_) denotes the Spearman rank correlation coefficient between attribute *k* and latent dimension *j*, and the absolute value captures both positive and negative monotonic relationships. For each property, the metric identifies the latent dimension with the strongest monotonic relationship (maximum absolute correlation). The Spearman correlation is distribution-free and captures monotonic relationships without assuming linearity, making it particularly suitable for evaluating the monotonic alignment enforced by attribute regularization.
- **Modularity** assesses whether each latent dimension depends on at most one attribute, measuring the degree to which individual dimensions are specialized for specific attributes. For each latent dimension *j*, mutual information with all attributes is computed, and the dimension is evaluated based on how exclusively it captures a single attribute:

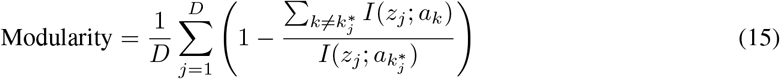

where *D* is the number of latent dimensions, *I*(*z*_*j*_; *a*_*k*_) denotes mutual information between latent dimension *j* and attribute *k*, and 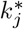 is the index of the attribute with highest mutual information with dimension *j*.
- **Mutual Information Gap (MIG)** evaluates disentanglement by measuring how much a single latent dimension dominates in capturing information about each attribute:

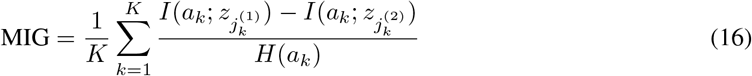

where *K* is the number of attributes, *I*(*a*_*k*_; *z*_*j*_) denotes mutual information between attribute *k* and latent dimension 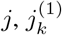 and 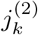 are the indices of latent dimensions with the highest and second-highest mutual information with attribute *k*, and *H*(*a*_*k*_) is the entropy of attribute *k*. Higher MIG scores indicate better disentanglement, where each attribute is primarily encoded in a distinct latent dimension rather than distributed across multiple dimensions.
- **Separated Attribute Predictability (SAP)** measures disentanglement by evaluating how well individual latent dimensions can predict attributes independently. For each attribute *k*, a classifier is trained on each latent dimension *j* separately to predict the attribute value, yielding prediction scores *S*_*k,j*_:

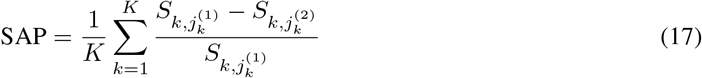

where *K* is the number of attributes, 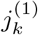 and 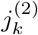 are the indices of the two most predictive latent dimensions for attribute *k* (ranked by prediction score), and *S*_*k,j*_ represents the prediction score (e.g., coefficient of determination *R*^2^ or classification accuracy) when predicting attribute *k* using only latent dimension *j*.

Results across all metrics are reported in Section A.7.

#### A.6.2 Sequence Quality

##### Fréchet Biological Distance

To evaluate the quality of generated peptides in biologically relevant embedding space, we compute Fréchet Biological Distance (FBD) using fine-tuned ESM2 embeddings. The ESM2-t12 model (Lin et al., 2023) was fine-tuned for binary antimicrobial activity classification using active/inactive labels with thresholds of ≤ 32 *µ*g/ml for active and ≥ 128 *µ*g/ml for inactive peptides.

FBD is computed analogously to Fréchet Inception Distance (Heusel et al., 2017) by modeling the distributions of ESM2 embeddings as multivariate Gaussians:

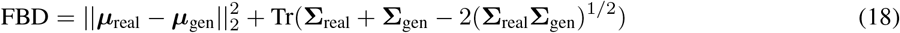

where ***µ*** and **Σ** represent the mean and covariance of the embedding distributions for real and generated peptides, respectively.

*FBD*_*train*_ measures how well generated peptides conform to the training distribution by computing FBD between generated sequences and the training dataset using ESM2 embeddings, being a proxy for peptide validity.

**Diversity** is a fraction of unique generated sequences not present in the training dataset.

**Novelty** is computed using average pairwise Levenshtein distance among generated sequences.

##### Fitness Score

The Fitness Score (Li et al., 2024) serves as a proxy for peptide amphiphilicity, a critical structural property for antimicrobial activity. Amphiphilicity, characterized by the spatial separation of hydrophobic and hydrophilic residues, enables peptides to interact with and disrupt bacterial membranes.

Notably, while the Fitness Score is not explicitly regularized during training, it is influenced indirectly through properties that we do control: charge and hydrophobicity, which directly affect the spatial distribution of polar and non-polar residues, and antimicrobial activity predictions, which implicitly capture amphiphilic requirements for membrane disruption. This emergent control demonstrates PepGlider’s ability to modulate complex structural properties through regularization of correlated physicochemical properties.

Let 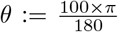 (the equivalent of 100 degrees in radians), and let both *h* and *hx* denote amino-acid scales. For a peptide sequence of length *L* with amino acids indexed by *i*, the Fitness Score is computed as:

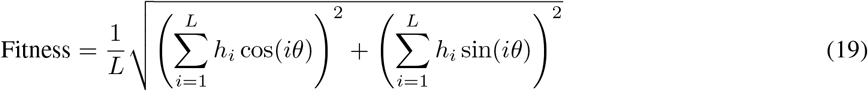

where *h*_*i*_ represents the hydrophobicity scale value for amino acid at position *i*. The angular parameter *θ* corresponds to an ideal *α*-helical geometry, and the summation captures the vectorial distribution of hydrophobic moments along the helical axis. Higher scores indicate greater amphiphilic character, reflecting optimal segregation of hydrophobic and hydrophilic residues necessary for membrane interaction and antimicrobial function.

**Supplementary Table S2:** Detailed property-specific disentanglement quality metrics results across physicochemical properties for PepGlider and baseline and ablation methods. Higher scores indicate better disentanglement for all metrics.

| Model | Property | Interpretability (↑) | SCC (↑) | Modularity (↑) | MIG (↑) | SAP (↑) |
| --- | --- | --- | --- | --- | --- | --- |
| VAE | Length | 0.542 | 0.754 | 0.893 | 0.012 | 0.022 |
|  | Charge | 0.088 | 0.320 | 0.660 | 0.001 | 0.026 |
|  | Hydrophobicity | 0.080 | 0.309 | 0.736 | 0.000 | 0.009 |
| HydrAMP | Length | 0.951 | 0.973 | 0.957 | 0.056 | 0.020 |
|  | Charge | 0.024 | 0.439 | 0.868 | 0.003 | 0.042 |
|  | Hydrophobicity | 0.178 | 0.440 | 0.867 | 0.000 | 0.029 |
| Transformer | Length | 0.014 | 0.289 | 0.799 | 0.005 | 0.038 |
|  | Charge | 0.022 | 0.333 | 0.723 | 0.004 | 0.013 |
|  | Hydrophobicity | 0.261 | 0.494 | 0.975 | 0.008 | 0.121 |
| ARVAE | Length | 0.994 | 0.995 | **0.999** | 0.895 | 0.948 |
|  | Charge | 0.947 | 0.994 | 0.979 | 0.280 | 0.655 |
|  | Hydrophobicity | 0.921 | 0.996 | 0.177 | 0.211 | 0.621 |
| **PepGlider** | Length | 0.977 | 0.995 | **0.999** | 0.856 | 0.927 |
|  | Charge | 0.934 | 0.991 | 0.980 | 0.292 | 0.649 |
|  | Hydrophobicity | 0.881 | **0.998** | 0.976 | 0.211 | 0.587 |
| PepGlider w/o normalization | Length | 0.994 | 0.995 | **0.999** | 0.884 | 0.953 |
|  | Charge | 0.958 | 0.992 | 0.981 | 0.297 | 0.670 |
|  | Hydrophobicity | 0.990 | 0.997 | 0.983 | 0.257 | 0.690 |
| PepGlider with signum | Length | 0.994 | 0.995 | **0.999** | **0.896** | 0.951 |
|  | Charge | 0.950 | 0.994 | 0.980 | 0.285 | 0.655 |
|  | Hydrophobicity | 0.921 | 0.996 | 0.973 | 0.177 | 0.612 |
| PepGlider w/ *z-score* normalization | Length | **0.996** | 0.995 | **0.999** | 0.882 | **0.959** |
|  | Charge | 0.966 | 0.982 | 0.982 | **0.321** | 0.668 |
|  | Hydrophobicity | 0.937 | **0.998** | 0.980 | 0.231 | 0.631 |

##### Biological Activity

We evaluate antimicrobial activity using APEX (Wan et al., 2024) predictions for *E. coli* and *S. aureus* providing species-specific MIC predictions for clinically relevant pathogens, as described in A.1.2. Toxicity assessment is performed using our trained hemolytic toxicity classifier as described in A.1.3

*FBD*_*active*_ evaluates similarity to experimentally validated antimicrobial peptides by computing FBD to a curated set of active peptides with documented MIC ≤ 32 *µ*g/ml against at least one bacterial strain in DBAASP, serving as antimicrobial potency proxy.

**Supplementary Table S3:** Detailed property-specific disentanglement quality metrics results across antimicrobial properties (MIC *E. coli*, MIC *S. aureus*, non-toxicity) for PepGlider and baseline methods. Higher scores indicate better disentanglement for all metrics.

| Model | Property | Interpretability (↑) | SCC (↑) | Modularity (↑) | MIG (↑) | SAP (↑) |
| --- | --- | --- | --- | --- | --- | --- |
| VAE | MIC *E. coli* | 0.087 | 0.287 | 0.815 | 0.001 | 0.029 |
|  | MIC *S. aureus* | 0.076 | 0.275 | 0.795 | 0.001 | 0.027 |
|  | Nontoxicity | 0.054 | 0.550 | 0.897 | 0.002 | 0.020 |
| HydrAMP | MIC *E. coli* | 0.000 | 0.295 | 0.928 | 0.000 | 0.020 |
|  | MIC *S. aureus* | 0.000 | 0.248 | 0.931 | 0.001 | 0.012 |
|  | Nontoxicity | 0.007 | 0.320 | 0.931 | 0.002 | 0.009 |
| Transformer | MIC *E. coli* | 0.129 | 0.365 | 0.945 | 0.001 | 0.008 |
|  | MIC *S. aureus* | 0.090 | 0.304 | 0.932 | 0.000 | 0.006 |
|  | Nontoxicity | 0.039 | 0.371 | 0.945 | 0.001 | 0.027 |
| **PepGlider** | MIC *E. coli* | 0.566 | 0.962 | 0.992 | 0.074 | 0.022 |
|  | MIC *S. aureus* | **0.758** | 0.933 | 0.991 | 0.062 | **0.236** |
|  | Nontoxicity | 0.332 | **0.992** | **0.994** | **0.075** | 0.123 |

### A.7 Extended Results

**Supplementary Figure S3:**
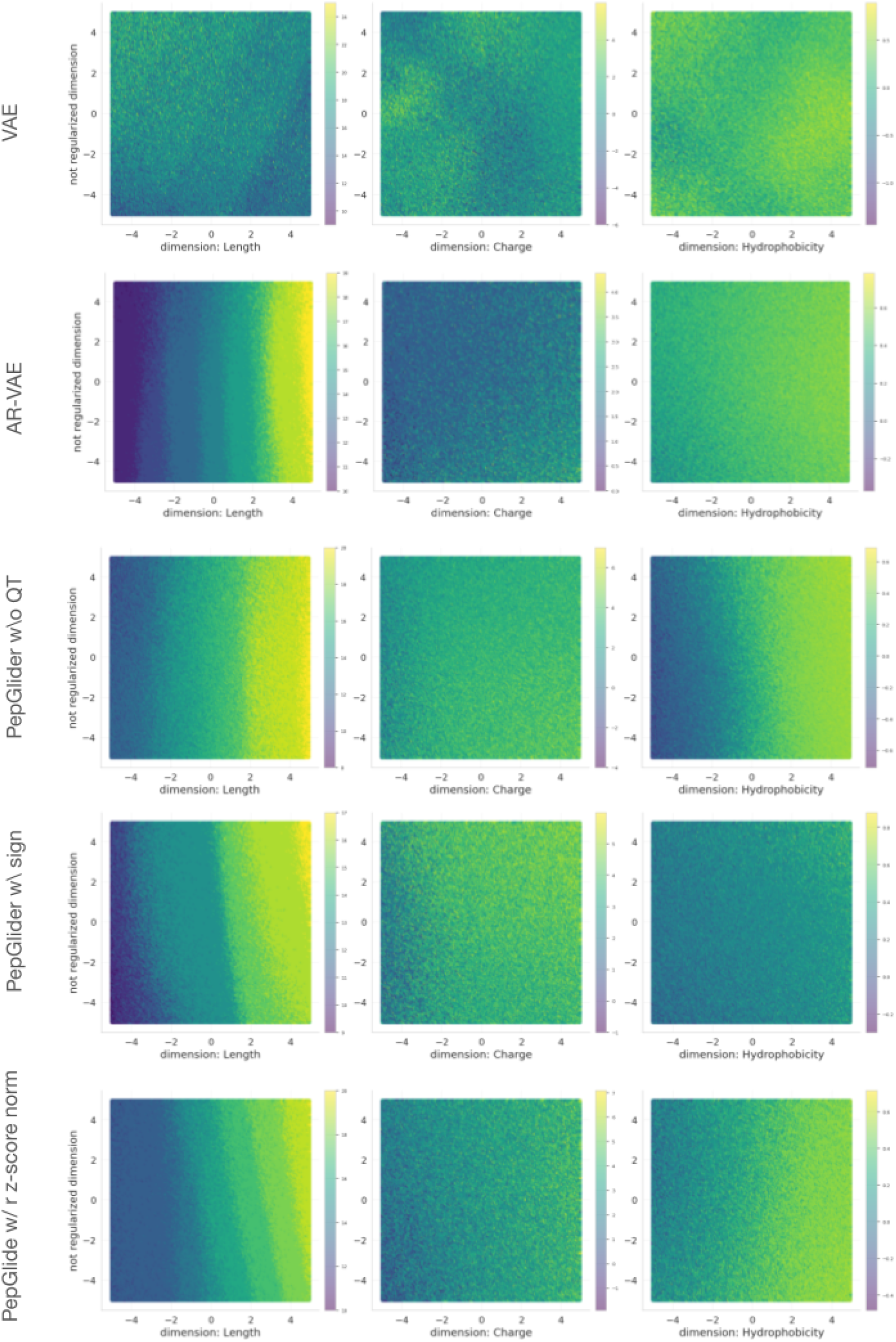
Latent space traversal. 2D property surface plots showing property transitions for length (left column), charge (middle column), and hydrophobicity (right column) across different model architectures. From top to bottom: VAE, AR-VAE, PepGlider without quantile transformation (w/o QT), PepGlider with signum (w/ sign), and PepGlider with z-score normalization (w/ z-score norm).

**Supplementary Figure S4:**
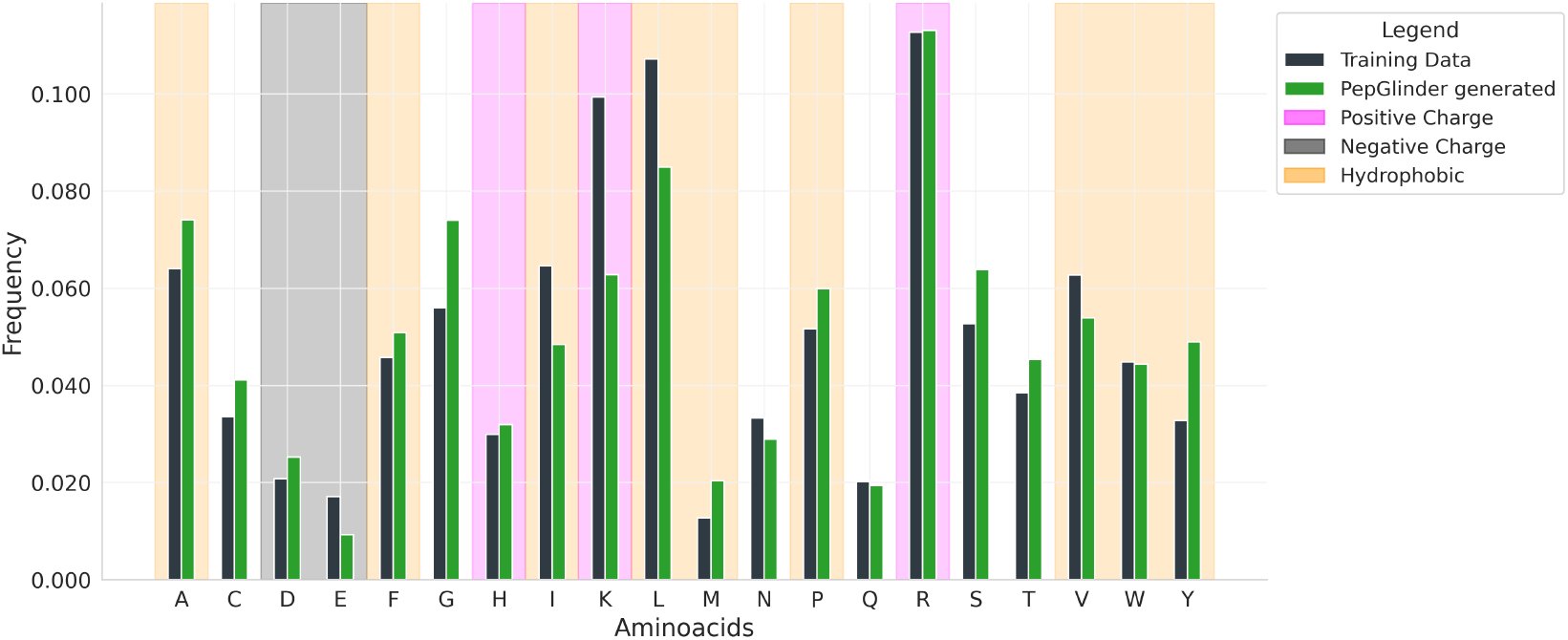
Reconstruction quality assessment for PepGlider-generated peptides. Amino acid frequency distributions comparing generated peptides (green bars) with training data (black bars). Amino acids crucial for hydrophobicity are highlighted in orange, and amino acids contributing to positive charge and negative charge are highlighted in pink and gray, respectively.

**Supplementary Figure S5:**
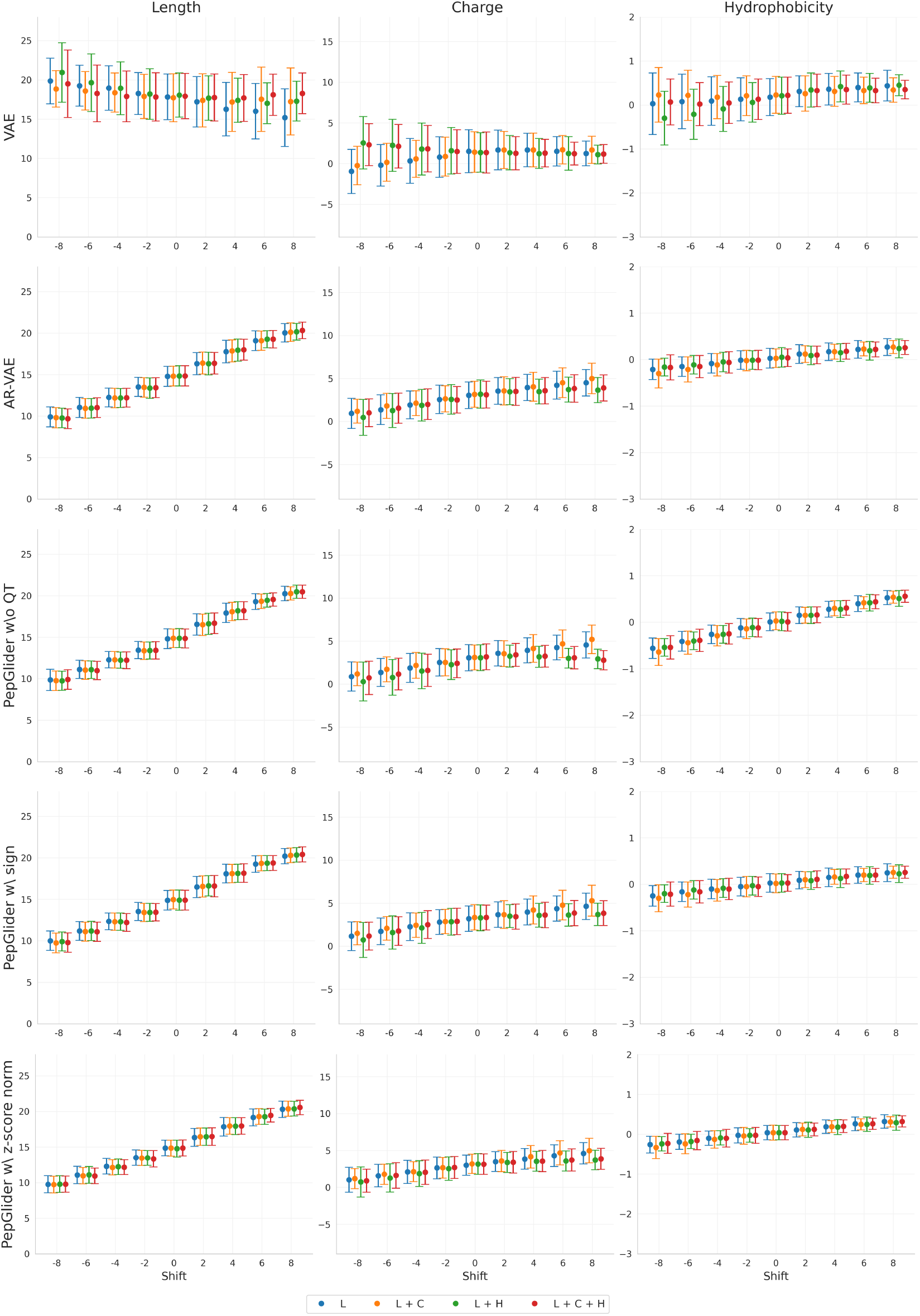
Multi-propoerty conditioning across model variants. Property values across latent space displacements for length (left column), charge (middle column), and hydrophobicity (right column) under single-property (L, C, or H), dual-property (L+C, L+H, or C+H), and tri-property (L+C+H) conditioning scenarios. Rows show different model architectures: VAE, AR-VAE, PepGlider without quantile transformation (w/o QT), PepGlider with signum (w/ sign), and PepGlider with z-score normalization (w/ z-score norm). Error bars represent mean ± standard deviation.

**Supplementary Figure S6:**
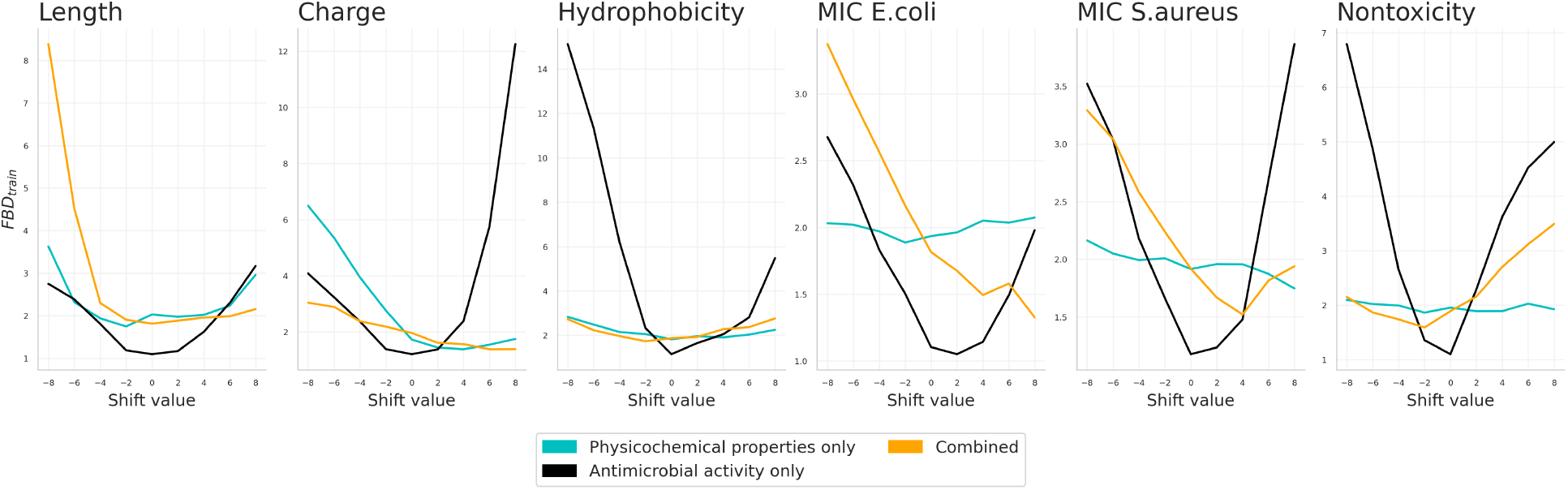
Validity across models with different sets of properties and regularized dimensions. *FBD*_*train*_ across latent shifts for models with different regularized property sets: physicochemical properties only (cyan), antimicrobial activity only (black), and combined physicochemical and activity properties (orange). Each panel shows validity when manipulating the corresponding property dimension across displacement.

**Supplementary Figure S7:**
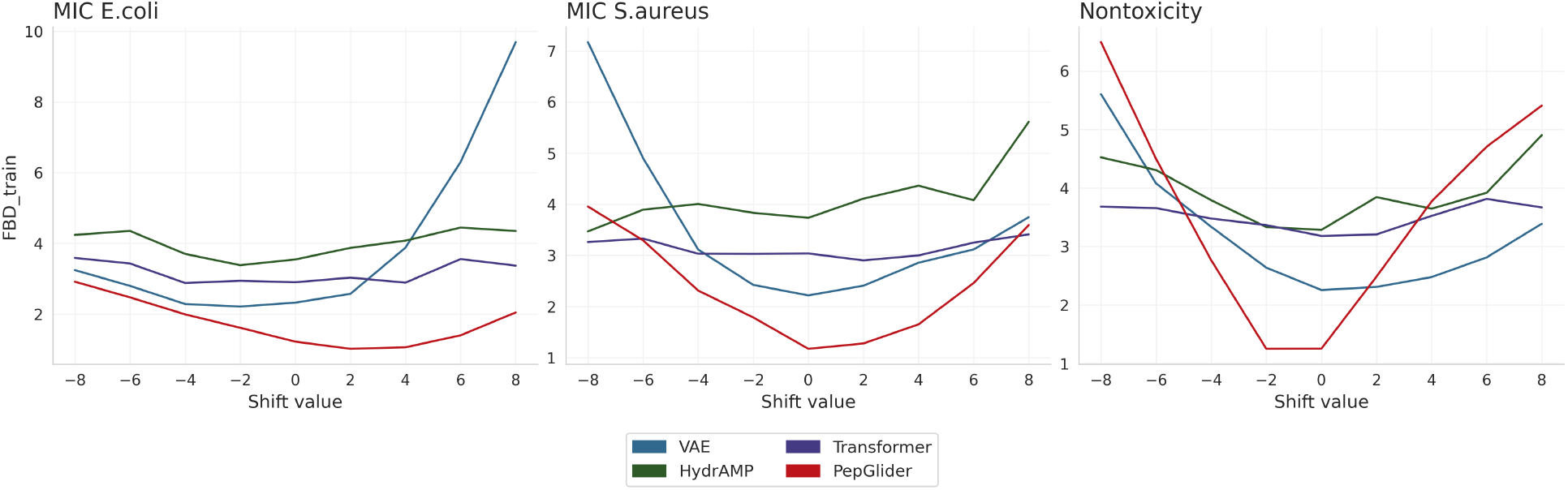
Model robustness to distribution shifts across antimicrobial activity metrics. *FBD*_*train*_ values for four generative models (VAE, Transformer, HydrAMP, PepGlider) evaluated on *E. coli* activity (APEX prediced MIC), *S. aureus* activity (APEX predited MIC), and non-toxicity across varying shift magnitudes.

